# Ccp1 depletion disrupts network integration of hippocampal parvalbumin interneurons

**DOI:** 10.1101/2024.11.08.622673

**Authors:** Romain Le Bail, Bernard Lakaye, Ira Espuny-Camacho, Carsten Janke, Maria M. Magiera, Dominique Engel, Carla G. Silva, Laurent Nguyen

## Abstract

Post-translational modifications (PTMs) of microtubules (MTs) endow them with specific properties that are essential for key cellular functions, such as axonal transport. Polyglutamylation, a PTM that accumulates in long-lived MTs, has been linked to neurodegeneration in the cerebellum when in excess. While hyperglutamylation of MTs leads to neurodegeneration and disrupts the function of specific neuronal subtypes like Purkinje cells, cortical neurons, and hippocampal excitatory neurons, little is known about its impact on inhibitory interneurons and their functional integration into local networks. In this study, we generated a conditional knockout mouse model to deplete cytosolic carboxypeptidase 1 (Ccp1) in GABAergic neurons, a key MT deglutamylase expressed by hippocampal interneurons. Our findings reveal that the loss of Ccp1 has a profound effect on hippocampal parvalbumin (*PV*)-expressing interneurons, impairing their axonal transport and reducing their perisomatic inhibition of pyramidal cells (PCs) in the CA2 region of the hippocampus.

**Research Topics:** Molecular Neuroscience, Cell Biology

**Highlights:**

- Different subtypes of hippocampal interneurons express unique sets of (de)glutamylases and show varying levels of protein posttranslational glutamylation.
- Parvalbumin interneurons become hyperglutamylated when Ccp1 activity is lost.
- The loss of Ccp1 disrupts axonal transport in interneurons and is associated with decreased perisomatic inhibition of hippocampal pyramidal cells in the CA2 region.

## Introduction

Microtubules (MTs) are polarized cytoskeletal elements essential for the establishment of neuronal morphology, and function as "tracks" for the intracellular transport of cargoes. This process is crucial, for example, for synapse formation and function^1,2^. In neurons, MTs are heterogeneous, with distinct molecular characteristics and tailored to undertake specific cellular roles^3^. This diversity could arise from the dynamic incorporation of different tubulin isotypes, the accumulation of various post-translational modifications (PTMs), and/or their association with different pools of microtubule-associated proteins (MAPs)^3^. The combination of tubulin isotypes and PTMs forms the foundation of the "tubulin code"^4^. As neurons differentiate and mature, they form long-lived MTs that accumulate various PTMs, including polyglutamylation, which involves the addition of glutamate residues to the γ-carboxyl group of gene-encoded glutamate on tubulins^5^. The generation of these glutamic acid side chains is catalyzed by members of the tubulin tyrosine ligase-like (Ttll) family of enzymes^6^. MT glutamylation is reversible, with the breakdown of glutamate side chains and branching points controlled by enzymes from the cytosolic carboxypeptidase (Ccp) family^6,7^. In neurons, glutamylation levels are tightly regulated, and MT hyperglutamylation has been shown to be causative of neurodegeneration in the *pcd* mouse model. This mouse line is characterized by a loss-of-function (LOF) mutation in the *Agtpbp1* gene, which encodes the deglutamylase Ccp1. In this model, the progressive degeneration of Purkinje neurons can be prevented by knocking down Ttll1, which restores physiological MT glutamylation levels^8,9^. Neurodegeneration in the *pcd* mouse brain is neuron-type specific, indicating that intrinsic cell properties contribute to this phenotype. In fact, cortical neurons are spared in this model, unlike Purkinje neurons, likely because Ccp1 LOF function is compensated by the high expression and activity of Ccp6. Consistently, simultaneous depletion of Ccp1 and Ccp6 results in axonal transport defects and neurodegeneration in the cerebral cortex after five months^7,8^. The molecular mechanism linking MT hyperglutamylation to neurodegeneration in the *pcd* mouse remains unclear. However, increased polyglutamylation causes reduced axonal transport of various cargoes in hippocampal pyramidal cells (PCs), significantly affecting mitochondrial motility, fusion, and fission, which could further compromise cell survival^8,10,11^. Mouse models lacking both Ccp1 and either Ttll1 or Ttll7 (the most abundant glutamylases) exhibit distinct brain phenotypes, further indicating that different MT glutamylation patterns generated by specific enzymes have unique effects on axonal transport and neuronal survival^12^. Ttll7 LOF does not prevent Purkinje neuron degeneration neither restore mitochondrial transport in the *pcd* mouse, suggesting that Ttll1 activity is responsible for the pathological phenotypes^12^.

Distinct brain regions express specific subsets of enzymes that regulate MT PTMs, leading to regional patterns of MT polyglutamylation, which may underpin specific cellular functions^7^. To test this hypothesis, we examined the role of the core enzymes controlling (de)glutamylation—Ccp1, Ccp6, and Ttll1—in hippocampal interneurons. Our previous work demonstrated that cortical interneurons express a wide range of Ttlls (glutamylation “writers”) and Ccps (glutamylation “erasers”) starting in early development. Conditional Ccp1 LOF in migrating interneurons results in MT hyperglutamylation, accompanied by mild MT hyperstabilization in newly formed neurite branches^13^. However, the functional consequences of MT hyperglutamylation in mature interneurons and whether Ccp1 loss leads to their progressive death have not been explored.

In this study, we used a conditional knockout mouse model to investigate the long-term impact of Ccp1 depletion in hippocampal interneurons. Our results reveal that Ccp1 loss leads to protein hyperglutamylation in hippocampal interneurons during adulthood and that interneuron subtypes exhibit distinct basal expression levels of (de)glutamylation enzymes. Parvalbumin (*PV*) interneurons express higher levels of Ttll1, Ccp1, and Ccp6 compared to somatostatin (*SST*) interneurons. *PV^+^* interneurons lacking Ccp1 activity, but not *SST^+^* interneurons, are hyperglutamylated and show axonal transport defects. Additionally, we observed a reduction of inhibitory *PV^+^* interneuron synapses on the soma of pyramidal cells (PCs) in the hippocampal CA2 region, correlated with a reduction in miniature inhibitory postsynaptic currents (mIPSCs), indicating alterations in the *PV^+^* interneuron-dependent hippocampal networks.

Our findings demonstrate that cellular alterations caused by Ccp1 LOF are neural subtype-specific. This study also supports the idea that protein hyperglutamylation may lead to a spectrum of phenotypes, with neurodegeneration being the most severe. The ultimate outcome could depend on how different cell types are sensitive to MT perturbation and how they regulate key enzymes involved in neurodegeneration. The specific axonal transport and network defects we observed in *PV^+^* hippocampal interneurons broaden the range of cell types and phenotypes affected by hyperglutamylation and highlight the importance of studying the tubulin code at a single-cell level.

## Results

### Differential expression of enzymes involved in glutamylation homeostasis in hippocampal interneuron subtypes

The level of MT glutamylation in neurons depends on the combined action of enzymes that act as writers and erasers of glutamylation^12^. However, the expression of these enzymes at the level of individual cells has been overlooked, especially in different types of hippocampal interneurons. To investigate the cellular distribution of single mRNA molecules, we used basescope on brain slices from three-month-old mice to detect and quantify the mRNAs of the main (de)glutamylases: Ccp1, Ccp6, and Ttll1. Our focus was on both *PV^+^* and *SST^+^* interneuron subtypes, which together make up most of the hippocampal interneurons (Figures 1A-1I)^14^. *PV^+^* interneurons are primarily found in or near the PC layer (*stratum pyramidale*) of the CA1, CA2, and CA3 regions, targeting PC somas to provide perisomatic inhibition^15^. In contrast, *SST^+^* interneurons are functionally diverse and spread across different layers of the hippocampus^16^. We detected mRNA molecules coding for Ccp1 (Figure 1B), Ccp6 (Figure 1E), and Ttll1 (Figure 1H) in both *PV* and *SST* interneurons in three-month-old mice. However, there were more mRNA puncta for Ccp1, Ccp6, and Ttll1 in *PV^+^* interneurons compared to *SST^+^* interneurons (Figures 1C, 1E, 1H). Overall, our findings indicate that both *PV^+^* and *SST^+^* interneurons in the hippocampus express the same key (de)glutamylation enzymes, but *PV^+^* interneurons show higher basal expression levels. This suggests that the function of *PV^+^* interneurons may require a tighter control of their MT (de)glutamylation levels, and a loss of Ccp1 would have a greater impact on the glutamylation balance in *PV^+^* interneurons compared to *SST^+^* interneurons.

**Figure 1.**
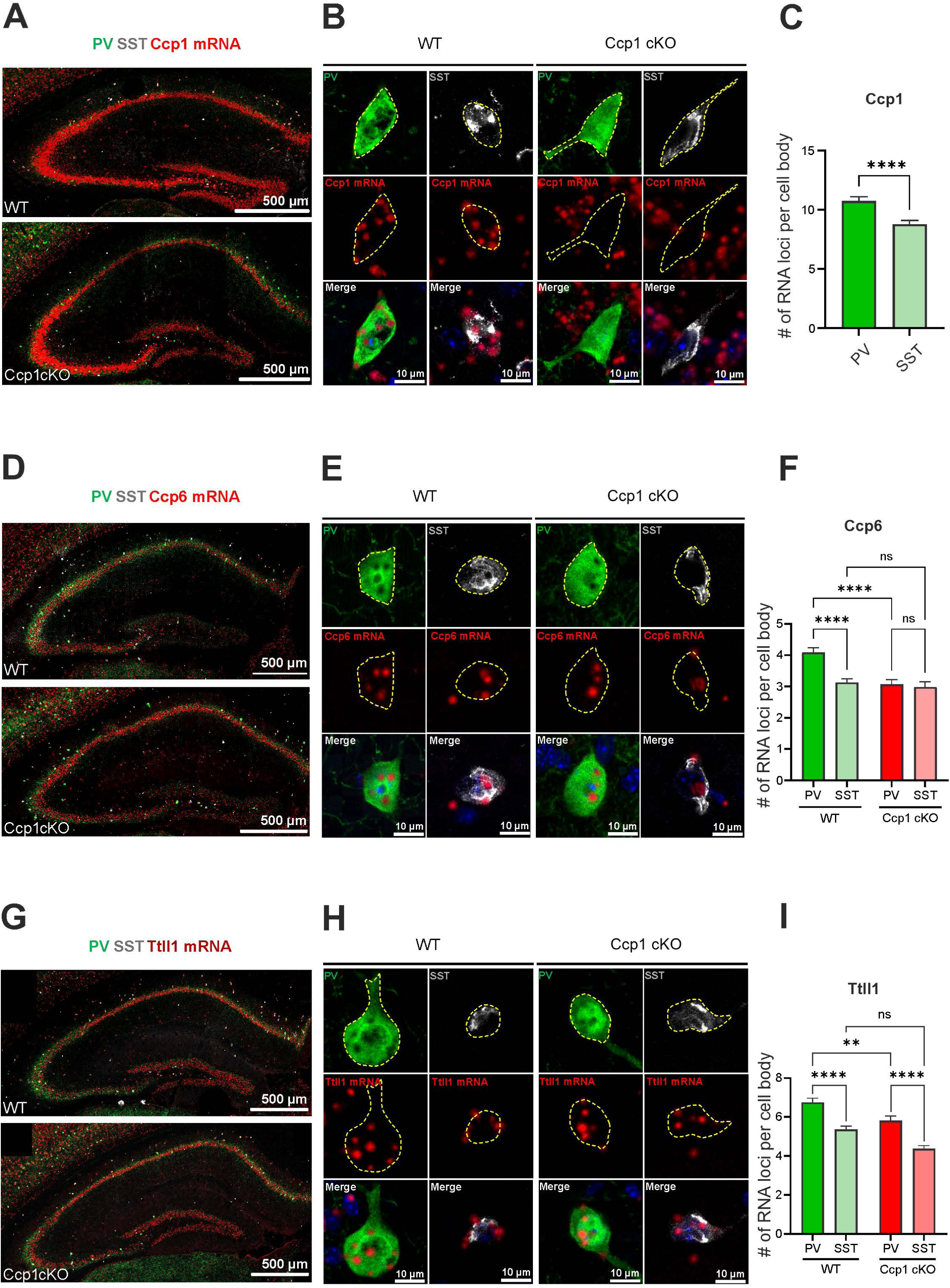
Differential expression of (de)glutamylation enzymes in SST^+^ and PV^+^ interneurons. (**A**, **D,** and **G**) Immunolabeling of hippocampi in brain slices from 3 months WT and Ccp1 cKO brains, showing mRNA expression (red) of *Ccp1*, *Ccp6*, and *Ttll1* together with *PV* (green) and *SST* (light grey) proteins. (**B**, **E**, and **H**) Immunolabeling of hippocampal *PV*^+^ and *SST*^+^ interneurons showing representative signal (red) for *Ccp1*, *Ccp6* and *Ttll1* mRNAs. Yellow doted lines delineate cell contours. (**C**) Quantification of the number of Ccp1 mRNA dots within the cell bodies of WT *PV*^+^ and *SST*^+^ interneurons. Because no signal is present for Ccp1 mRNA in Ccp1 cKO interneurons, quantification was only performed in WT animals. *PV*^+^ WT = 156 neurons, *SST*^+^ WT = 165 neurons from 3 WT mice. For statistical analysis, a Mann-Whitney analysis was conducted (p < 0,0001). (**F**,**I**) Quantification of the number of *Ccp6* or *Ttll1* mRNA dots within cell bodies of WT and Ccp1 cKO *PV*^+^ and *SST*+ interneurons. For *Ccp6* mRNA, *PV*^+^ WT = 266 neurons, *SST*^+^ WT = 255 neurons, *PV*^+^ cKO = 171 neurons, *SST*^+^ cKO = 107 neurons from 4 WT and 3 Ccp1 cKO mice. For *Ttll1* mRNA, *PV*-WT = 225 neurons, *SST*^+^ WT = 234 neurons, *PV*^+^ cKO = 196 neurons, *SST*^+^ cKO = 201 neurons from 4 WT and 3 Ccp1 cKO mice. For statistical analysis, data were transformed according to the formula Y = sqrt(Y+1) to obtain a normal distribution, then a two-way ANOVA analysis was conducted. ** = p < 0.1, *** = p < 0.01, **** = p< 0.001.

### Loss of Ccp1 alters the expression of genes involved in glutamylation homeostasis in PV^+^ interneurons

We subsequently examined whether the conditional deletion of *Ccp1* in hippocampal interneurons induces compensatory alterations in the expression of *Ccp6* and *Ttll1*. To address this, we utilized the conditional mouse model *Dlx5,6 Cre-GFP; Ccp1^f/f^* (hereafter referred to as *Ccp1 cKO*), in which *Ccp1* is selectively inactivated in GABAergic forebrain neurons during development^13,17^. Given that the *Dlx5,6* enhancer is inactive in adulthood, we crossed *Ccp1 cKO* mice with *ROSA-Lox-STOP-Lox-YFP* mice to permanently label GABAergic neurons. The total number of hippocampal interneurons in three-month-old mice was consistent across genotypes (Figures S1A-S1B). These findings suggest that the loss of *Ccp1* activity does not result in interneuron degeneration, contrary to previous reports on Purkinje neurons in the cerebellum^18^. *Ccp6* is strongly expressed in neurons, and its modulation has been shown to compensate for *Ccp1* LOF in certain neuronal populations^7,8^. Additionally, *Ttll1* expression is downregulated in certain cell types to prevent MT hyperglutamylation and cell death. We therefore examined how *PV^+^*and *SST^+^* interneurons regulate *Ccp6* and *Ttll1* expression following the conditional loss of *Ccp1* in *Ccp1 cKO* mice. Quantification of *Ccp6* and *Ttll1* mRNA loci in hippocampal *Ccp1 cKO PV^+^* and *SST^+^* interneurons revealed a reduction in the expression of both *Ccp6* and *Ttll1* in *PV^+^*interneurons, but not in *SST^+^* interneurons (Figures 1F, 1I). While a reduction in *Ttll1* expression may partially mitigate polyglutamylation, the reduction in *Ccp6* expression suggests that *PV* interneurons fail to compensate for *Ccp1* loss through *Ccp6* upregulation. These findings indicate that *PV^+^* and *SST^+^* interneurons possess distinct intrinsic capacities to regulate protein glutamylation in response to *Ccp1* loss, and imply a failure of *PV* interneurons to upregulate deglutamylases as compensation for the loss of *Ccp1*.

### PV^+^ interneurons accumulate hyperglutamylated proteins upon loss of Ccp1

Immunolabeling of brain slices from three-month-old WT and *Ccp1 cKO* mice was conducted to detect hippocampal *PV^+^*and *SST^+^* interneurons, along with polyglutamylated proteins (C-terminally located linear glutamate chains consisting of at least four glutamic acid residues; polyE^19^) (Figures 2A-2B). We observed that protein glutamylation was specifically abundant in *PV* but not *SST* hippocampal interneurons in *Ccp1 cKO* brains (Figure 2C). We then investigated whether *PV* interneurons located in the three main hippocampal regions (CA1, CA2, and CA3) exhibited hyperglutamylation following *Ccp1* loss of activity. To delineate these anatomical hippocampal regions, we performed immunolabeling for *RGS14*, a regulator of G-protein signalling that is enriched in CA2 dendritic spines (Figure S2A, blue staining)^20^. We observed that *PV* interneurons exhibited uniform levels of glutamylation across hippocampal regions (Figure S2B). These findings suggest that the loss of *Ccp1* expression induces protein hyperglutamylation in *PV^+^* interneurons distributed throughout the hippocampus, but not in *SST* interneurons.

**Figure 2.**
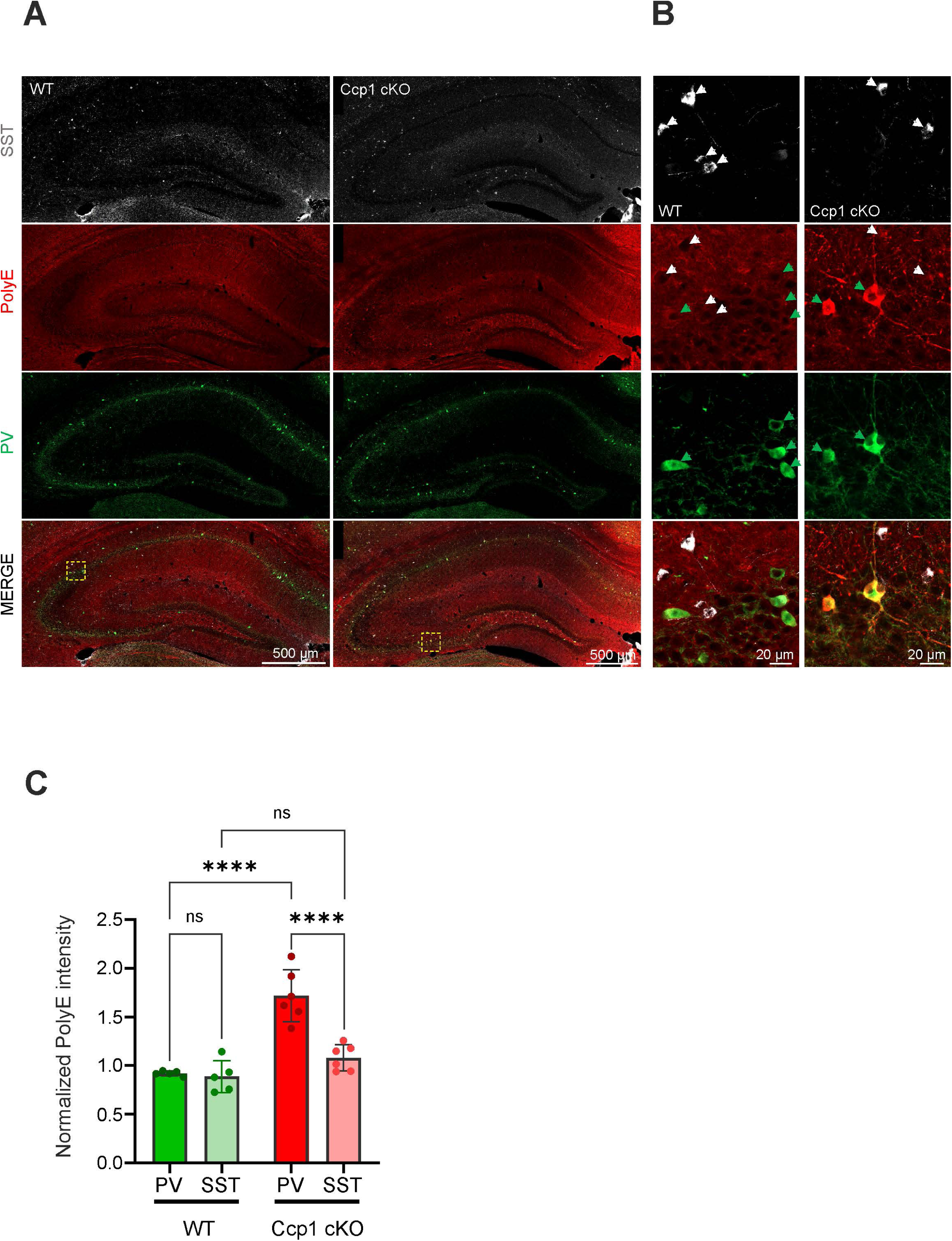
PV^+^ interneurons are hyperglutamylated in Ccp1 cKO adult hippocampi. **(A)** Representative images of hippocampi immunolabeled with *SST* (light grey), PolyE (red), and *PV* (green) in WT and Ccp1 cKO three months old mice. PolyE^+^ labeling strongly enriched in *PV*^+^ interneurons (green arrowheads) in the Ccp1 cKO hippocampus. Yellow dotted square corresponds to the close up area in panel B. (**B**) Close up area of representative *PV*^+^ and *SST*^+^ interneurons immunolabeled to detect PolyE in the hippocampus of WT and Ccp1 cKO mice. White and green arrows point to the cell bodies of *SST* and *PV* interneurons, respectively. In Ccp1 cKO, *PV*^+^ but not *SST*^+^ interneurons show strong PolyE immunoreactivity. (**C**) Average PolyE intensity in *PV*^+^ and *SST*^+^ interneurons of WT and Ccp1 cKO mice. PolyE staining intensity was normalized to the intensity of the surrounding hippocampal tissue. *PV*^+^ interneurons show significant hyperglutamylation in Ccp1 cKO. Each dot on the graph represents the average PolyE intensity for all *PV*^+^ or *SST*^+^ interneurons analyzed in one animal. WT: n=5 CCP1cKO n= 6. A two-way ANOVA was conducted after confirmation of normal data distribution, **** = p < 0.0001

The primary protein targets for (de)glutamylation are alpha and beta tubulins, and MT hyperglutamylation has previously been linked to defects in axonal transport across various cargo types^8,10^. Since *SST* interneurons did not exhibit hyperglutamylation following *Ccp1* loss, we tested whether *PV* interneurons displayed axonal transport defects. Given that protein hyperglutamylation levels were consistent across all *PV* interneurons (Figure S2B), we cultured hippocampal cells from E17.5 mouse embryos to measure and compare axonal lysosome transport in interneurons that either expressed or lacked *Ccp1* at DIV 15 (Figure S3A). Hippocampal cultures from both WT and *Ccp1 cKO* mice included predominantly excitatory neurons (*NeuN^+^, GFP^-^*) and a smaller fraction of interneurons (*NeuN^+^, GFP^+^*) (Figures S3A-S3C). Among the interneurons (*GFP^+^*), only a small fraction expressed *PV* (ranging from 10% to 14%; Figures S3D-S3E). Culturing cells at low density on gridded dishes enabled time-lapse imaging of acidic vesicles in interneuron neurites, followed by the identification of cell type through *PV* immunohistochemistry (Figure S3F). This approach allowed for the visualization of interneuron neurites and subsequent molecular transport analysis. To monitor lysosome transport, we labelled lysosomes with Lysotracker and analyzed their movement along the main proximal neurites using real-time video microscopy (Figures S4A-S4D; Supplemental movie S1). The largest proximal neurite was considered the developing axon of interneurons. We measured anterograde and retrograde lysosome transport and analyzed their speed, as well as the fraction of time the lysosomes were in motion or paused during transport. No differences were observed in the average speed of anterograde or retrograde lysosome transport between genotypes (Figures 3A-3B, 3F-3G). However, the fraction of time that lysosomes spent moving in *PV^+^* interneurons was reduced following the loss of *Ccp1* (Figure 3C), while the fraction of time lysosomes were paused was significantly increased in *Ccp1 cKO PV*^+^ interneurons compared to their controls (Figure 3D). Additionally, the fraction of immotile lysosomes—defined as particles that exhibited no motion throughout the entire duration of the recording—was also higher in *Ccp1 cKO PV^+^* interneurons (Figure 3E). Comparable analyses were conducted in *PV^-^* interneurons, where no differences were detected in neither anterograde nor retrograde lysosomal transport (Figures S4E-S4F, S4J-S4K). While the proportion of lysosomes moving per unit of time in *PV^-^* interneurons was consistent across genotypes, the fraction of time that lysosomes were paused was increased following the loss of *Ccp1* (Figures S4C, S4G, S4J-S4K). However, the proportion of immotile lysosomes remained unchanged (Figure S4I). Overall, our results confirm that the loss of *Ccp1* activity induces significant changes in MT-dependent transport dynamics, which correlates with hyperglutamylation in hippocampal *PV^+^* interneurons. Given that axonal transport defects in neurons have previously been associated with neurodegeneration, we quantified the numbers of both *SST^+^* and *PV^+^* interneurons and found no differences between three-month-old WT and *Ccp1 cKO* mice (Figures 4A-4D).

**Figure 3.**
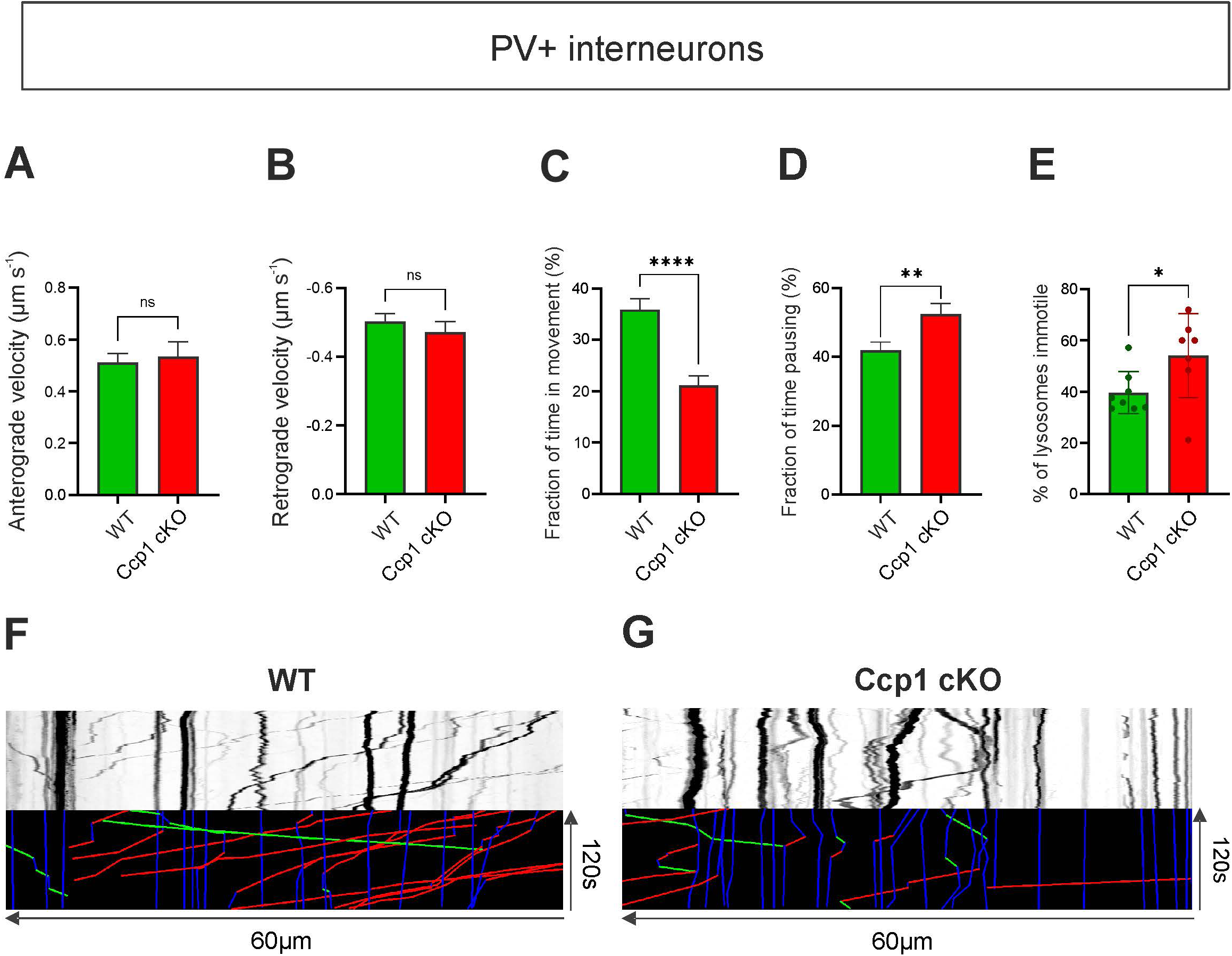
Vesicular transport is perturbed in Ccp1 cKO PV^+^ interneuron neurites. **(A)** Histograms of the anterograde velocity of lysosome (lysotracker) trafficking in hippocampal *PV*^+^ interneurons, mouse genotypes are indicated on the histogram. Only the fractions of movement when lysosomes travel away from the cell body are considered. When lysosomes are moving in the anterograde direction, no change in velocity was detected in Ccp1 cKO *PV*^+^ interneurons. The number of lysosomes with anterograde movement in this analysis is 99 for WT and 79 for Ccp1 cKO. A Mann-Whitney test was conducted (p = 0.3803). (**B**) Histograms of the retrograde velocity of lysosome (lysotracker) trafficking in hippocampal *PV*^+^ interneurons, mouse genotypes are indicated on the histogram. The retrograde velocity only considers the fractions of movement when lysosomes travel towards the cell body and no changes were detected in Ccp1 cKO *PV*^+^ interneurons. The number of lysososmes with retrograde movement in this analysis is 169 for WT and 110 for Ccp1 cKO. A Mann-Whitney test was conducted (p = 0.2339). (**C**) Histogram of the fraction of time lysosome spent moving in *PV*^+^ interneuron from both mouse genotypes. This analysis takes into account all lysosomes tracked, including motile and stationary ones and measures the percentage of time in movement for each lysosome. A threshold of 0.1 µm/s was established below which lysosomes are considered immotile. Lysosomes are less motile in the Ccp1 cKO condition as compared to WT. All lysosomes that were tracked were included in this analysis for a total of 342 and 334 ones in WT and Ccp1 cKO, respectively. A Mann-Whitney test was conducted (p < 0.0001). (**D**) Histogram of the fraction of time motile lysosomes pose in *PV*^+^ interneurons from both mouse genotypes. This measurement only considers motile lysosomes for at least a portion of the movie and represents the percentage of time spent with a speed < 0.1 µm/s, which is considered as pausing. Motile lysosomes spend significantly more time pausing in Ccp1 cKO *PV*^+^ interneurons. A total of 211 and 148 motile lysosomes were included for the WT and Ccp1 cKO conditions, respectively. A Mann-Whitney test was conducted (p = 0.006). (**E**) Histogram of the percentage of immotile lysosome sin *PV*^+^ intereneurons. Lysosomes are considered as immotile if their instantaneous speed is < 0.1µm/s for the whole duration of the movie. For this analysis, the proportion of immotile lysosomes in each neuron was calculated. Individual values on the graph represent single neurons. *PV*^+^ interneurons show a higher fraction of immotile lysosomes in the Ccp1 cKO condition. Eight and seven *PV*^+^ interneurons were recorded in the WT and Ccp1 cKO conditions, respectively, and out of 3 unrelated experiments. A Mann-Whitney test was conducted (p = 0.0370). (**F**-**G**) Kymographs of moving lysosomes for each genotype in *PV*^+^ interneurons. Colored kymographs represent the tracks that were manually traced where blue lines correspond to time pausing, green lines anterograde movement and red lines retrograde movement.

**Figure 4.**
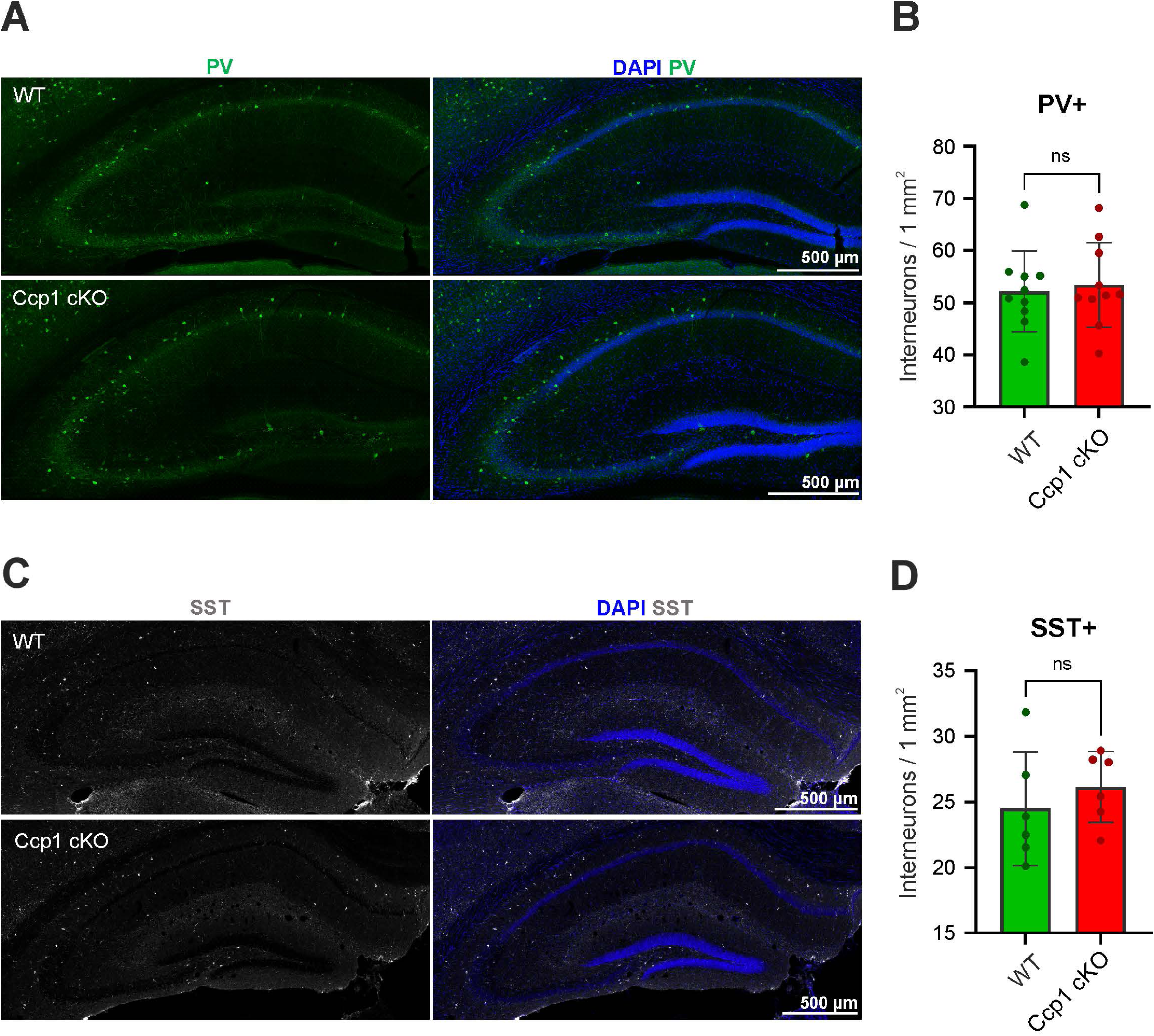
CCP1 loss does not trigger degeneration of adult hippocampal interneurons. **(A)** Immunolabeling of WT and hippocampi from 3 months old WT and Ccp1 cKO mice, showing the distribution of *PV*^+^ interneurons (green) and a nuclei counterstaining with Dapi (blue). (**B**) Histogram of the density of *PV*^+^ interneurons in the CA1, CA2 and CA3 regions normalized by the surface of the analyzed area. n =10 mice for both conditions, an unpaired t-student test was performed (p = 0.7256). (**C**) Immunolabeling of WT and hippocampi from 3 months old WT and Ccp1 cKO mice, showing the distribution of *SST*^+^ interneurons (light grey) and a nuclei counterstaining with Dapi (blue). **D**) Histogram of tdensity of *SST*^+^ interneurons in the CA1, CA2 and CA3 regions normalized by the surface of the analyzed area. n =6 mice for both conditions, an unpaired t-student test was performed (p = 0.4438).

While the loss of *Ccp1* did not trigger degeneration of *PV^+^*interneurons in the hippocampus at three months of age, it may still impair their physiological function and network integration. *PV^+^* interneurons primarily target the soma and proximal dendrites of PCs. To investigate this, we performed immunohistochemistry to label GABAergic synapses in the hippocampal formation. We used antibodies against the vesicular GABA transporter (VGAT)^21^ to label pre-synaptic inhibitory synapses and gephyrin^22^ to label postsynaptic sites (Figure 5A). Synaptic contacts were analyzed using super-resolution microscopy and defined as puncta where both VGAT and gephyrin were co-labeled (supplemental Movie S2)^23^. We first examined the CA1, CA2, and CA3 regions and their respective strata (*stratum pyramidale* (SP), *stratum oriens* (SO), and *stratum radiatum/lacunosum moleculare* (SLM)). No changes in synapse numbers were observed in the SP, SO, or SLM of the CA1 and CA3 regions (Figures S5A-S5C, 5F-5H). However, we detected a reduced number of synapses in the CA2 SP (Figure 5B) and SO regions in *Ccp1 cKO* hippocampi (Figures S5D), corresponding to the strata where *PV^+^* interneurons predominantly project^24^. We then confirmed in vitro that the loss of Ccp1 activity in interneurons resulted in a reduction of perisomatic inhibition of pyramidal cells (PCs), which is primarily mediated by *PV+* basket cells (Figures 5C, 5D, 5E). We observed fewer gephyrin-VGAT puncta above the somas of PCs that were targeted by multiple GABAergic terminals. A reduced number of perisomatic inhibitory boutons is expected to lead to a reduction in miniature inhibitory postsynaptic currents (mIPSCs). These synaptic events correspond to the spontaneous release of one quantum of GABA^25^. We performed patch-clamp recordings on PCs from the CA2 region, identifiable by an expansion of the PC layer thickness and confirmed *post hoc* by the localization of the recorded neuron, labeled with biocytin, in a region immunoreactive for *RGS14*, corresponding to CA2 (Figure 5F). We recorded mIPSCs in PCs from WT and *Ccp1 cKO* brain slices (Figures 5G, 5J) and quantified the cumulative probability of the peak amplitude (Figure 5H) and inter-event interval (Figure 5K) of mIPSCs. We found that the average peak amplitude of mIPSCs was unchanged in the *Ccp1 cKO* condition (Figure 5I), while the frequency was significantly decreased (Figure 5L). These results support the anatomical data indicating a reduction in the number of GABAergic synapses onto PCs and, consequently, a decrease in the number of GABA release sites in interneurons from the *Ccp1 cKO*. In contrast to mIPSCs, the average amplitude of spontaneous IPSCs (sIPSCs) was slightly, but not significantly, increased in PCs from *Ccp1 cKO* mice compared to those from WT mice (Figure S5K). However, the frequency of sIPSCs remained unchanged (Figure S5N). Perisomatic inhibitory cells (aka basket cells) are known to regulate the firing of pyramidal cells^26^. We therefore examined the passive and active properties of CA2 PCs in both WT and *Ccp1 cKO* animals. We found no significant differences between the genotypes (Figures S5I, S5O, S5M, S5S; Table 1). We did observe however a trend towards lower membrane potential (Figure S5Q), higher input resistance (Figure S5R), and increased number of action potentials (Figure S5S) in CA2 PCs from *Ccp1 cKO* mice, as well as a tendency for a higher firing rate in response to current injection (Figure S5M). Although the passive and active properties of CA2 PCs were not significantly altered, the reduced perisomatic inhibition could lead to a slight increase in the excitability of these PCs. Together, our results suggest that perisomatic inhibition of PCs is impaired, through a reduction in mIPSCs, particularly in the CA2 region of the hippocampus. These findings indicate that some local circuit alterations in the hippocampus may occur as a consequence of *Ccp1* loss in PV^+^ interneurons, independently of neurodegeneration.

**Figure 5.**
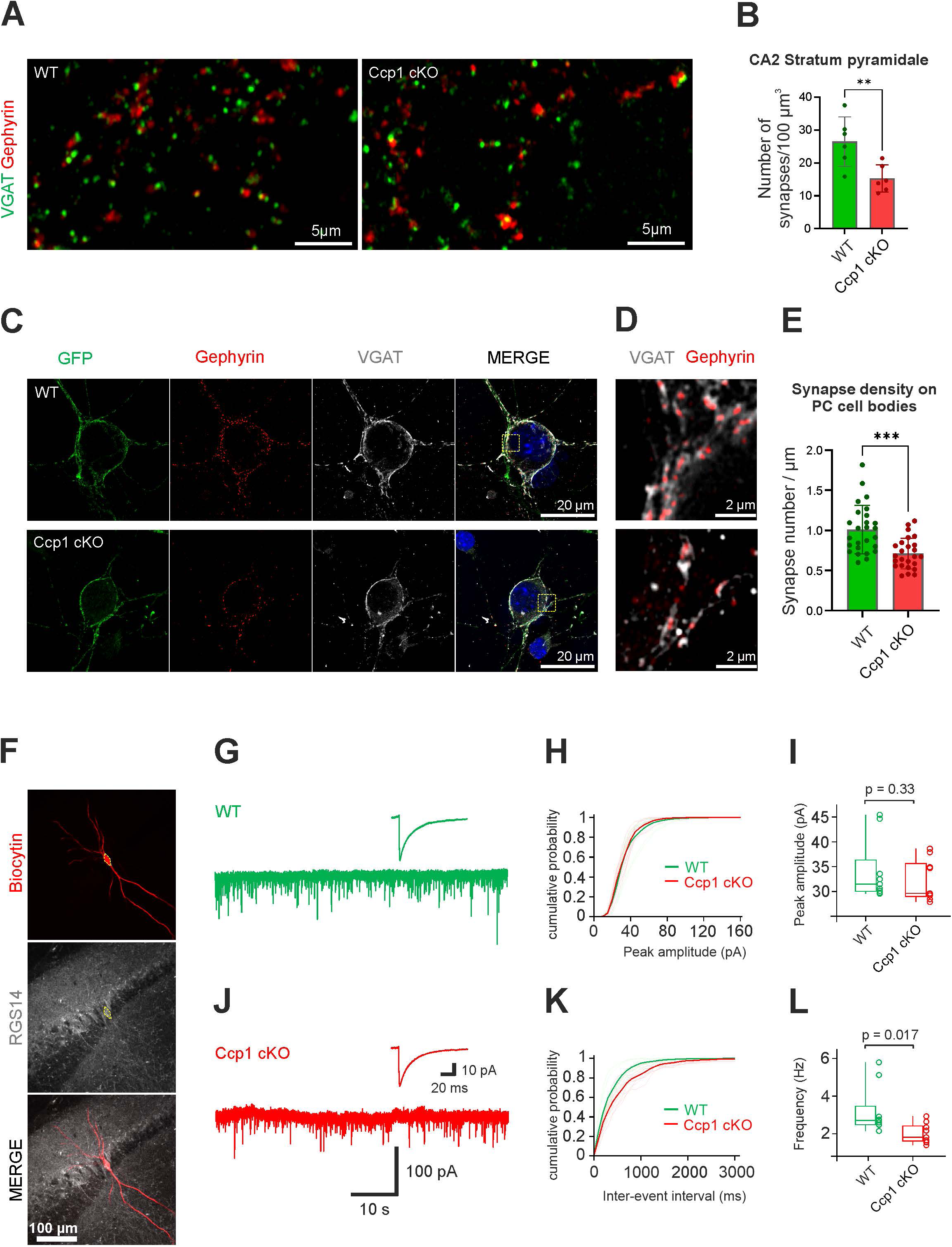
CCP1 loss results in decrease of inhibitory synapses in the CA2 stratum pyramidale of the hippocampus. **(A)** Immunolabeling of synapses where pre and postsynaptic markers are shown VGAT (green) and gephyrin (red), respectively. This analysis was performed on 6 WT and 6 Ccp1 cKO 3 month-old mice, an unpaired t-student test was conducted (p = 0.0093). (**B**) Quantification of inhibitory synapses density in the CA2 *stratum pyramidale*. The colocalization of one gephyrin red) puncta and one VGAT (green) puncta was considered as an inhibitory synapse. This analysis was performed on 6 WT and 6 Ccp1 cKO 3 month-old mice, an unpaired t-student test was conducted (p = 0.0093). (**C**) Representative images of WT or Ccp1 cKO neurons innervated by GFP^+^ interneuron axons. Immunolabeling detects VGAT (light grey) and gephyrin (red). Only GFP^-^ neurons (principal cells, or PC) whose cell body was innervated by GFP^+^ fibers were considered in this analysis. (**D**) Magnification of the yellow dotted square in panel (**C**) showing the colocalization of VGAT with Gephyrin punctae. (**E**) Histogram of density of inhibitory synapses along GFP^+^ fibers innervating the cell body of PC. The number of VGAT; gephyrin puncta that colocalize was normalized to the length of GFP^+^ fibers that innervate the cell body. A total of 26 and 25 cell bodies were analyzed in WT and CCP1 cKO respectively, from 3 independent experiments. An unpaired t-student test was conducted (p < 0.0001). (**F**) Representative image of a CA2 PC injected with biocytin in a sagittal hippocampal slice from a 2 month-old mouse. The yellow dotted line outlines the cell body of the biocytin labelled cell to highlight the colocalization with the CA2 PCs marker RGS14. (**G**, **J**) Representative traces of miniature IPSCs (mIPSCs) in a CA2 PC at -70 mV from a WT slice (**G**, green trace) and from a Ccp1 cKO slice (**J**, red trace). External solution contained 1 mM TTX, 20 mM D-AP5, and 10 mM CNQX to block action potentials and excitatory synaptic activity. Inset (**G**) shows average mIPSC at expanded timescale (average from 1,778 [9 WT neurons] single events and inset (**J**) average from 1,104 [9 cKO neurons] single events). (**H**) Cumulative histograms of mIPSC amplitude (light green: from 9 individual PCs of WT mice; thick green: average of WT data; light red: from 9 individual PCs of CCP1 cKO mice; red: average of CCP1 cKO data). (**I**) Boxplots of mIPSC peak amplitude. Data from 9 WT neurons and 9 cKO neurons. Averaged mIPSC amplitude was similar in control and cKO neurons (Mann-Whitney test, p=0.33). (**K**) Cumulative histograms of mIPSC inter-event interval (light green: from 9 individual PCs of WT mice; thick green: average of WT data; light red: from individual PCs of CCP1 cKO mice; red: average of CCP1 cKO data). (**L**) Boxplots of mIPSC frequency. Data from 9 WT neurons and 9 cKO neurons. Averaged mIPSC frequency was significantly decreased in CCP1 cKO neurons when comparing to control neurons (Mann-Whitney test, p=0.017). In box plots (**I** and **L**), horizontal lines represent median; boxes, quartiles; whiskers, extreme data points; and single points, data from individual experiments.

## Discussion

In the present study, we examined the consequences of enhanced polyglutamylation resulting from the Ccp1 LOF specifically in GABAergic interneurons, with a focus on axonal transport, synaptic density, and synaptic release. We delineated the expression profiles of *Ccp1*, *Ccp6*, and *Ttll1* mRNA in mature hippocampal *PV^+^* and *SST^+^* interneurons, and found no compensatory changes in the expression of *Ccp6* and *Ttll1* in the absence of Ccp1 specifically in *PV*^+^ interneurons. Moreover, we observed a significant increase in polyglutamylation intensity in *PV*^+^ interneurons compared to *SST*^+^ interneurons. This cell type-specific change was associated with an alteration of axonal transport in *PV*^+^ interneurons, while only mild disruptions were detected in *SST*^+^ interneurons. Finally, we demonstrated a reduction in the density of perisomatic GABAergic synapses and inhibitory synaptic transmission in pyramidal cells (PCs) within the CA2 region of the hippocampus.

In the absence of Ccp1, protein hyperglutamylation was specifically observed in *PV^+^* GABAergic interneurons, while no such changes were noted in *SST^+^* cells. This enhanced polyglutamylation may be attributed to upstream modifications in the mRNA expression levels of glutamylases and deglutamylases. In WT animals, higher basal levels of *Ccp1*, *Ccp6*, and *Ttll1* mRNA were quantified in *PV*^+^ cells compared to *SST*^+^ interneurons. In the absence of Ccp1, *Ccp6* mRNA and *Ttll1* mRNA levels were reduced in *PV*^+^ cells but not in *SST*^+^ cells. Therefore, *PV*^+^ cells exhibit particular sensitivity to the loss of Ccp1, with protein hyperglutamylation primarily resulting from the absence of the originally high levels of *Ccp1* mRNA expression, as *Ttl1* mRNA was only moderately downregulated and *Ccp6* mRNA exhibited a decrease in expression levels. The altered balance of polyglutamylation and deglutamylation specifically in *PV*^+^ interneurons indicates a cell-type-specific regulation of polyglutamylation. This observation underscores the necessity of investigating the effects of Ccp1 loss at the cellular level, as specific neuronal subpopulations may be uniquely affected. Indeed, among cerebellar GABAergic neurons, Purkinje cells exhibit a pronounced dependence on the presence of Ccp1, as its loss leads to protein hyperglutamylation and rapid neurodegeneration of Purkinje neurons^7^. These severe consequences may be attributable to the very low expression of *Ccp6* and the absence of other members of the *Ccp* family that could compensate for protein deglutamylation. Analysis of lysosomal transport in the neurites of hippocampal Ccp1 cKO *PV*^+^ and *PV*^-^ interneurons in vitro revealed significant transport defects in *PV*^+^ interneurons, which closely resembled those previously described for hippocampal neurons^8,10^. Specifically, Ccp1 cKO *PV*^+^ interneurons exhibited a marked reduction in overall vesicle motility, characterized by increased pausing time and a higher proportion of stationary vesicles; however, the anterograde and retrograde velocities remained unchanged. This suggests that MT hyperglutamylation may disrupt the initiation of cargo movement or induce motor inactivity or detachment. Conversely, Ccp1 cKO *PV*^-^ interneurons demonstrated only an increase in the pausing time of motile vesicles, with no significant change in the proportion of stationary vesicles. Given that polyglutamylation levels are elevated in *Ccp1* cKO *PV*^+^ interneurons, it is plausible that lysosomes remain stationary for extended periods, thereby increasing the overall proportion of vesicles that are immobile throughout the recording duration. The phenotype observed in Ccp1 cKO *PV*^-^ interneurons may reflect a milder alteration in transport, albeit governed by similar mechanisms as those in *Ccp1* cKO *PV*^+^ interneurons. Notably, Ccp1 cKO *PV*^-^ interneurons did not exhibit increased polyglutamylation levels *in vivo* at three months of age. Similar to cortical neurons, it is possible that glutamylation levels are increased during the perinatal period due to the lack of compensation by other *Ccps*, which may account for the mild transport defects observed in adult *PV*^-^ interneurons^7,13^.

The absence of Ccp1 led to a decrease in the density of inhibitory synapses specifically in the *stratum pyramidale* (SP) and *stratum oriens* (SO) of the CA2 region in the adult hippocampus, accompanied by a reduction in the frequency of miniature inhibitory postsynaptic currents (mIPSCs). Inhibitory synapses were identified based on the colocalization of presynaptic vesicular GABA transporter (VGAT) and postsynaptic gephyrin, indicating that all GABAergic synapses were considered regardless of the specific identity of the presynaptic GABAergic cell^23^. In the CA2 SP and SO, the majority of GABAergic boutons originate from *PV*^+^ basket cells, whose somata are located in the SP^27^. Consequently, the reduction in inhibitory synapses is primarily attributed to a decrease in presynaptic GABAergic boutons from *PV*^+^ interneurons that are hyperglutamylated. The decrease in mIPSC frequency observed in Ccp1 cKO mice supports the reduction of GABAergic synapses. Miniature currents are synaptic currents detected at the soma because they originate from synapses located in or in dendrites close to the soma^28,29^, while spontaneous currents occurring further out in the dendrites remain undetectable at the soma^30^. Therefore, mIPSCs in CA2 pyramidal cells are predominantly produced by perisomatic *PV*^+^ inhibitory neurons, including basket and chandelier cells^29^. Notably, the frequency of mIPSCs, but not spontaneous IPSCs (sIPSCs), was decreased in pyramidal cells in the absence of Ccp1. The lack of a decrease in sIPSC frequency may be attributable to a compensatory increase in GABA release by other interneurons^31^. The trend toward larger sIPSC amplitudes in Ccp1 cKO mice might indicate an additional compensatory mechanism for the reduced perisomatic inhibition. The reduction in synaptic release associated with altered MT polyglutamylation levels has been previously documented. The absence of functional polyglutamylase PGs1, a subunit of alpha-tubulin-selective polyglutamylase, results in a reduction of MT polyglutamylation and a mis-localization of the kinesin Kif1A in neurites. This leads to a decreased density of docked vesicles at presynaptic terminals, ultimately resulting in a diminished release during sustained synaptic transmission^32^. Increased levels of polyglutamylation resulting from the loss of spastin, a MT-severing protein, have been shown to reduce both glutamatergic synapses and the frequency of miniature excitatory postsynaptic currents (mEPSCs)^33^. This reduction in the number of synaptic vesicles in presynaptic boutons has been hypothesized as a potential causal link to the decreased frequency of mEPSCs, stemming from abnormal delivery and distribution of synaptic cargoes^33^. Physiological levels of polyglutamylation are therefore essential for maintaining axonal transport and, consequently, ensuring sufficient accumulation of docked vesicles at presynaptic terminals, which is necessary for both basal and high-frequency synaptic vesicular release. The observed reduction in synapses may also result from axonal degeneration. In the absence of both Ccp1 and Ccp6, axons in the cerebral cortex exhibit signs of degeneration, including axonal lysis and accumulation of cellular organelles, potentially caused by defects in axonal transport^8^. Disturbed axonal transport is a common feature observed in several neurodegenerative diseases^34^. GABAergic innervation by local *PV*^+^ interneurons is particularly prominent in the hippocampal CA2 region^27^. These interneurons play a crucial role in mediating strong feedforward inhibition in CA2 upon the excitation of Schaffer collaterals^35^. A reduction in the frequency of mIPSCs, likely due to a decrease in the density of perisomatic inhibitory synapses, could result in an increased excitation/inhibition ratio, thereby diminishing feedforward inhibition. Changes in inhibitory synaptic transmission associated with dysfunctional *PV*^+^ interneurons have been observed in several neurodevelopmental disorders, including autism spectrum disorders^36^, epilepsy^37^, fragile X syndrome^38^, and Rett syndrome^39^.

In this study, we observed that the expression levels of Ccp1, Ttll1, and Ccp6 mRNA are increased in wild-type (WT) *PV*^+^ interneurons compared to wild-type *SST*^+^ interneurons. Additional experiments investigating the expression levels of other Ttll and Ccp genes, as well as their respective activities, may provide further insights into why *PV*^+^ interneurons, but not *SST^+^* interneurons, exhibit hyperglutamylation in the adult mouse hippocampus. Despite the hyperglutamylation of *PV*^+^ interneurons observed at three months in Ccp1 conditional knockout (cKO) mice, we did not find evidence of neurodegeneration. Future assessments of GABAergic interneuron populations at later time points could clarify whether the hyperglutamylation of *PV*^+^ interneurons ultimately leads to degeneration, as it has been reported in the cortex at five months^8^. The synaptic defects identified in Ccp1 cKO mice may result in functional and behavioral alterations. Given that the loss of synapses is primarily localized to the CA2 region of the hippocampus, it is plausible to hypothesize that social memory may be impaired in Ccp1 cKO mice^40^. A comprehensive electrophysiological analysis could be conducted to probe the functionality of inhibitory microcircuits, possibly through the stimulation of Schaffer collaterals. *PV*^+^ interneurons receive robust afferent inputs from CA3 pyramidal cells^41^, which subsequently inhibit CA2 pyramidal cells via feedforward inhibition^42^. Additionally, a behavioral screening approach to assess spatial and social memory in Ccp1 cKO mice could be implemented to determine whether potential defects in microcircuitry lead to broader alterations in memory encoding.

Altogether, our data suggest that disrupting the precise and sometimes cell-specific regulation of protein glutamylation homeostasis may have a broader impact on the establishment and/or maintenance of neuronal circuits.

## Material and methods

### Animals

All animals were maintained under standard conditions, including a 12-hour light/12-hour dark cycle, with free access to food and water, and a stable environment with a temperature of 19-22°C and humidity of 40-50%. All animal procedures complied with the guidelines of the Belgian Ministry of Agriculture and adhered to the European Community Laboratory Animal Care and Use Regulations (86/609/CEE, Journal Official des Communautés Européennes L358, 18 December 1986). The animal experiments detailed in this chapter received ethical approval from the University of Liège’s ethics committee under protocols #1912, #2177, and #2455.

The study utilized male mice of the CD1 genetic background, sourced from Janvier Labs (Saint-Berthevin, France). Transgenic lines, including Dlx5,6 CRE-GFP, Ccp1 lox/lox, and ROSA-FloxStop YFP, were backcrossed to the CD1 background before being used in experiments ^13^. To avoid inbreeding, transgenic colonies were periodically bred with CD1 mice from Janvier Labs. Wild-type (WT) mice refer to male Dlx5,6 CRE-GFP/+ mice, while Ccp1 conditional knockout (cKO) mice are male Dlx5,6 CRE-GFP/+; Ccp1 lox/lox mice. In the Ccp1 cKO mice, exons 20 and 21 of the Ccp1 gene are specifically excised in the Dlx5,6 lineage through CRE-mediated recombination, targeting interneurons (INs) in the cortex and hippocampus. WT mice serve as controls for CRE and GFP expression in the Dlx5,6 lineage, maintaining intact Ccp1 alleles. For interneuron quantification in the adult hippocampus, WT and Ccp1 cKO mice were crossed with homozygous ROSA-FloxStop YFP CD1 mice to label and examine interneuron expression at 3 months.

### Immunohistochemistry

For brain collection, mice were anesthetized with an intraperitoneal injection of pentobarbital (Euthasol, 150 mg/kg). Once fully unconscious, they were perfused intracardially with 4% paraformaldehyde (PFA) in phosphate-buffered saline (PBS: 137 mM NaCl, 2.7 mM KCl, 10 mM NaLHPOL, 1.76 mM KHLPOL, pH 7.4). The brains were then extracted and placed in 4% PFA at 4°C overnight. The next day, the brains were rinsed with PBS and cryoprotected overnight at 4°C in 30% sucrose solution prepared in PBS. Cryoprotected brains were embedded in optimal cutting temperature (OCT) medium (Richard-Allan Scientific Neg50), frozen on dry ice, and stored at -80°C until further use.

For immunohistochemistry on brain tissue, 30 µm coronal sections were cut using a LEICA cryostat (LEICA – CM30505) and transferred into PBS. In certain experiments, antigen retrieval was carried out with citrate buffer (Agilent DAKO, S169984-2). Floating sections were placed in a sealed container with antigen retrieval solution and incubated in a water bath at 80°C for 25 minutes. Refer to Table 1 for details on specific experiments requiring antigen retrieval. The sections were then permeabilized and blocked in PBS containing 0.3% Triton X-100 (Sigma Aldrich, T8787) and 10% normal donkey serum (NDS, Jackson ImmunoResearch, RRID: AB_2337258) for 1 hour at room temperature (RT) under gentle agitation. Primary antibodies were diluted in antibody solution (PBS, 0.3% Triton X-100, 5% NDS), and the sections were incubated overnight at either 4°C or RT (see Table 1). Following three 15-minute washes with PBS containing 0.3% Triton X-100, the sections were incubated with DAPI (2 µg/mL, Sigma Aldrich, D9542) and appropriate secondary antibodies (Thermo Fisher Scientific or Jackson ImmunoResearch, 1:500) diluted in antibody solution for 2 hours at RT. This was followed by two washes in PBS with 0.3% Triton X-100 and a final wash in PBS. The sections were then mounted onto Superfrost slides (Fisher Scientific, Epredia J1800AMNZ) using pencil brushes. After 1 hour of drying, any excess salt was removed by a quick rinse in MilliQ water, and slides were covered with a glass coverslip using Fluoromount aqueous mounting medium (Sigma Aldrich, F4680). Preparation of samples for immunohistochemistry of synapses in vivo followed a protocol adapted from^43^.

Brains were rapidly extracted following euthanasia via intraperitoneal injection of pentobarbital (Euthasol) and immediately submerged in isopropanol chilled to -80°C. The brains were then stored at -80°C until cryosectioning. Coronal sections, 12 µm thick, were cut and collected onto Superfrost slides, which were promptly placed at -20°C to prevent degradation of the unfixed tissue. After cryosectioning, the slices were briefly fixed with 0.4% PFA in PBS using microwave irradiation (900W for 30 seconds). The slides were then washed with PBS, and the immunohistochemistry protocol outlined previously was applied for further processing.

For immunohistochemistry on primary hippocampal cultures, cells were grown on 12 mm coverslips, and the same protocol as described above was followed. Confocal images were acquired using the Airyscan mode on a Zeiss LSM 880 or Zeiss LSM 980 microscope (GIGA-Imaging Platform), depending on the experiment. All immunohistochemical analyses for a given experiment were performed concurrently and imaged using the same equipment and settings to ensure consistency. Image analysis was conducted with Fiji (ImageJ) or QuPath, depending on the specific requirements of the experiments.

### Semi-automated analysis of inhibitory synapses in hippocampal slices

Tile images of the CA1, CA2, and CA3 regions (as illustrated in Figure S6) were captured using a 63x oil objective on a Zeiss LSM880 microscope in Airyscan mode. The *stratum oriens* (SO), *stratum pyramidale* (SP), and *stratum radiatum* (SRL) regions were cropped as depicted in Figure S6. Semi-automated analysis of gephyrin puncta colocalizing with VGAT puncta was conducted using Imaris v9 software (Figure S7). Images were blinded to minimize experimenter bias during the manual analysis steps. Background subtraction (1 sigma) was applied to each channel, and gephyrin and VGAT puncta were identified using the spot detection feature. For each image, the estimated spot size was set to 400 nm, and a detection threshold was manually determined using the mean intensity filter. Inhibitory synapses were defined as gephyrin spots located within 400 nm of VGAT spots and filtered using the “shortest distance to spots” function.

### BaseScope

Brains were perfused and fixed overnight in 4% PFA as described in the “immunohistochemistry” paragraph. 12µm slices were cut at the cryostat, collected on superfrost slides and stored at -80°C before experiments. Custom probes were designed to detect Ccp1 (Advanced Cell Diagnostics, Cat. No. 1113561-C1), Ccp6 (Advanced Cell Diagnostics, Cat. No. 1143731-C1) and Ttll1 (Advanced Cell Diagnostics, Cat. No. 1158941-C1) mRNA using the BaseScope v2 detection kit REDTM (Advanced cell diagnostics, Cat. No. 323900) according to manufacturer instructions. Concomitant detection of PV and SST via immunohistochemistry was performed using the RNA-Protein Co-detection Ancillary kit (Advanced cell diagnostics, Cat. No. 323180). The Ccp1 probe was designed to hybridize only to exons 20 and 21 of Ccp1 which are removed in Ccp1 cKO animals therefore controlling for adequate excision of these exons in Ccp1 cKO INs.

Slides were first washed with PBS, baked for 30min at 60°C, post-fixed with 4% PFA for 15min at 4°C and the tissue was then dehydrated with successive ethanol baths (50%-70%-100%-100%) for 5min at room temperature. RNAscopeTM Hydrogen Peroxide was applied to the sections for 10min at RT and target retrieval was then performed by immersion of the slides in 100°C Co-detection Target Retrieval (Advanced Cell Diagnostics, Cat. No. 323180) for 5min. Slides were washed in water and once in PBS-T (PBS with 0,1% Tween-20). Slices were dried and primary antibodies diluted in Co-detection Antibody Diluent (Advanced Cell Diagnostics, Cat. No. 323180) were applied at 4°C overnight. The next day, slides were washed with PBS-T for 2 minutes twice and fixed with 4% PFA for 30min at RT. Additional PBS-T washes were performed and slices were treated with RNAscopeTM Protease III (Advanced Cell Diagnostics, Cat. No. 322337) for 30min at 40°C in an HybEZTM Oven (Advanced Cell Diagnostics, Cat. No. 310010). Slides were washed with distilled water and probes were hybridized for 2h, followed by serial amplifications (BaseScope v2 detection kit REDTM, Advanced cell diagnostics, Cat. No. 323900). Two washes with wash buffer reagent (Advanced Cell Diagnostics, Cat. No. 310091) were performed between each step. Probe hybridization and amplification steps 1-6 were all performed at 40°C in HybEZTM oven while amplification steps 7 and 8 were performed at RT. The signal was then revealed by incubating slides with BaseScopeTM Fast RED for 10min at RT. Finally, slides were wash buffer reagent and Co-Detection Blocker (Advanced Cell Diagnostics, Cat. No. 323180) was applied for 15min at 40°C and washed in wash buffer and PBS-T. DAPI (2µg/mL, Sigma, D9542) and secondary antibodies diluted 1:300 in Co-Detection Antibody Diluent were incubated for 1h at RT. Slides were washed in PBS-T several times and coverslips were mounted using FluoromountTM aqueous mounting media.

### Primary hippocampal cultures

Hippocampal cultures were prepared using a protocol adapted from^44^. For immunohistochemistry and axonal transport recordings, 12 mm glass coverslips in 24-well plates and 35 mm glass-bottom petri dishes (Mattek, P35G-1.5-14-CGRD-D) were used, respectively. Coverslips and dishes were coated by incubating with poly-ornithine (0.1 mg/mL in water, Sigma Aldrich #P4638) at 37°C for 45 minutes, followed by three rinses with water and overnight incubation with laminin (5 µg/mL in PBS, Sigma Aldrich #L2020) at 4°C. Before cell seeding, dishes were washed three times with PBS.

E17.5 hippocampi from WT or Ccp1 cKO mice were micro-dissected in Hank’s balanced saline solution (HBSS, Biowest, L0612-500), and the meninges were carefully removed. The hippocampi were enzymatically dissociated at 37°C for 25 minutes in a solution of 1:10 DNase (0.1%, Sigma, D5025) and 9:10 Papain (CellSystems, #LK003178) in HBSS, with the volume adjusted according to the number of hippocampi in the Eppendorf tube. After dissociation, the hippocampi were rinsed twice with 500 µL of Dulbecco’s modified Eagle medium (DMEM) supplemented with 10% fetal bovine serum (FBS) and 1:100 penicillin/streptomycin (Biowest, L0022-100). Mechanical dissociation was performed using a 1000 µL micropipette, and the cell suspension was filtered through a 40 µm mesh (Greiner, 542040).

The cells were counted, and 30,000 cells were seeded per well of 24-well plates or in glass-bottom petri dishes (Mattek, P35G-1.5-14-CGRD-D) for live imaging. Two hours after plating, complete neurobasal medium was added, consisting of Neurobasal Medium (Thermo Fisher, 12348017) supplemented with GlutaMAX (1:100, Thermo Fisher, 35050038), B-27 Supplement (1:50, Thermo Fisher, 17504044), and penicillin/streptomycin (1:100). Half of the medium was replaced with fresh complete neurobasal medium every 2-3 days. Cultures were maintained until DIV15, at which point they were either fixed in 4% PFA for 10 minutes at room temperature or used for neuronal transport recordings.

### Neuronal transport recording and analysis

Lysosomal transport in live neurons was recorded in WT and Ccp1 cKO hippocampal cultures at DIV15. In these cultures, GFP is expressed under the control of the Dlx5,6 enhancer to label INs in both WT and Ccp1 cKO mice. To specifically identify PV INs, neurons were cultured in glass-bottom dishes with an embedded grid (Mattek, P35G-1.5-14-CGRD-D) that allowed for precise positioning and imaging under the microscope (Figure S4A). Hippocampal cultures were maintained for 15 days as described earlier, and complete neurobasal medium without phenol red was used in the days leading up to imaging to minimize autofluorescence.

On the day of imaging, LysoTracker™ Red (100 nM, L7528) was added to the culture medium for 30 minutes at 37°C. Transport was recorded immediately using a 63x oil objective on a Zeiss LSM 980 confocal microscope, with environmental conditions maintained at 37°C and 5% COL. Lysosomal transport was imaged exclusively in the proximal neurites of GFP+ INs, with reference images taken in the green channel for each neuron, along with its grid position (Figure S4B). Movies were recorded for 2 minutes using Fast Airyscan mode with a frame rate of 600 ms to capture lysosome motility (Figure S4C). Multiple neurites from the same neuron were sometimes analyzed, with data collected from 25 neurons per genotype, across three independent cultures.

Immediately after imaging, the dishes were fixed in 4% PFA at room temperature for 10 minutes and washed three times with PBS. Immunohistochemistry for GFP and parvalbumin was performed using the previously described protocol (Figure S4D). The grid position and neuronal morphology from the reference images were used to locate GFP+ INs and determine their PV expression status. Neurons that could not be located after immunohistochemistry were excluded from the analysis due to unknown PV expression. The intensity of the PV signal within the soma was measured using Fiji to classify INs as either PV- or PV+. Lysosomal transport analysis was conducted with the KymoToolBox plugin^45^. Kymographs were generated for each movie, and lysosome paths were traced with segmented lines, capturing runs with direction and velocity changes. Summary data for each lysosome track were then extracted to determine anterograde and retrograde velocities, as well as pausing times.

### Slice preparation

Slices were prepared from the brains of 3 month-old swiss male mice. Animals were kept in an oxygenated chamber for >10 min before being anaesthetized with isoflurane (IsoFlo, Zoetis, Belgium; 4% added to the inspiration air flow). After decapitation, the brain was rapidly dissected out and immersed in ice-cold slicing solution containing: 87 NaCl, 25 NaHCO_3_, 10 D-glucose, 75 sucrose, 2.5 KCl, 1.25 NaH_2_PO_4_, 0.5 CaCl_2_ and 7 MgCl_2_, equilibrated with 95 % O_2_ / 5 % CO_2_ (pH 7.4, ∼325 mOsm) as previously described^46,47^. Sagittal 300-μm-thick slices were cut from the dorsal level of the hippocampus^48^ at 2-4 mm from midline using a vibratome (Leica VT-1200, Nussloch, Germany). Slices were stored in a reserve chamber containing slicing solution at 34°C for at least 30 min and subsequently at room temperature (20 – 25°C).

### Patch-clamp recording

Neurons were visualized using infrared IR-Dodt gradient contrast (IR-DGC) optics on a Zeiss FS microscope equipped with an IR camera (Newvicon tube in NC-70 Dage-MTI). In hippocampal slices, the CA2 region was visually identified and pyramidal cells (PCs) were selected in CA2 close to the transition of CA2 to CA1. The correct localization of the recorded PCs in CA2 was validated using post hoc immunohistochemistry of RGS14. For recordings, the slices were perfused with saline containing in mM: 125 NaCl, 25 NaHCO_3_, 25 D-glucose, 2.5 KCl, 1.25 NaH_2_PO_4_, 2 CaCl_2_ and 1 MgCl_2_ (equilibrated with 95% O_2_/5% CO_2_ gas mixture, pH 7.4, ∼315 mOsm) and maintained at room temperature. Patch pipettes were pulled from thick-walled borosilicate glass tubing (outer diameter: 2 mm, inner diameter: 1 mm; Hilgenberg, Germany) with a horizontal puller (P-97, Sutter Instruments). The composition of the internal solution was, in mM: 110 KCl, 30 K-gluconate, 2 MgCl_2_, 2 Na_2_ATP, 10 EGTA, 10 HEPES and 0.2% biocytin (pH = 7.2, osmolarity: ∼315 mOsm). Pipette resistance was 2.4-4.3 MΩ. Series resistance (6-12.5 MΩ) was not compensated in voltage-clamp, but carefully monitored during the experiments using 5 mV, 40 ms depolarizing test pulses. Synaptic currents were recorded in the voltage-clamp configuration with a holding potential of -70 mV. Neurons with a holding current larger than -130 pA were not included in the analysis. Miniature IPSCs were pharmacologically isolated in the presence of 10 μM 6-cyano-7-nitroquinoxaline-2,3-dione (CNQX), 20 μM D-2-amino-5-phosphonopentanoic acid (D-AP5) and 1 μM tetrodotoxin (TTX). Resting membrane potential was measured in current-clamp immediately after rupturing the membrane and entering the whole-cell mode. In current-clamp, a holding current was injected to maintain neurons at a membrane potential of ∼-70 mV. The input resistance was calculated from voltage responses to current pulses of 1 s duration (-50 pA to 50 pA amplitude, incremented in steps of 50 pA). Data points in the plot of the voltage measured 900–1000 msec after the onset of the pulse against the current step amplitude were fitted by linear regression. Sag measurements were obtained using a 1s long hyperpolarizing current (-200 pA). The average sag ratio was calculated using [(1 – ΔV_ss_ / ΔV_min_) × 100%], where ΔV_ss_ = RMP - V_ss_, ΔV_min_ = RMP - V_min_, RPM is the resting membrane potential, V_ss_ is the steady-state potential and V_min_ is the initial lowest potential (George et al., 2009). Recordings were performed at room temperature (20 – 25°C).

### Chemicals

Chemicals were as follows: tetrodotoxin citrate (TTX, Tocris) and biocytin (Life Technologies). The remaining chemicals were purchased from Sigma-Aldrich (Belgium).

### Data acquisition and analysis

Recordings were performed using an Axopatch 200B amplifier (Molecular devices, Palo Alto, CA) connected to a PC via a Digidata 1440A interface (Molecular devices). Data were acquired with pClamp 10.4 (Molecular devices). Capacitive currents were recorded in voltage-clamp and filtered at 10 kHz. Membrane potential and firing were examined in current clamp. Currents were low pass filtered (Bessel) at 10 kHz and acquired at 10 kHz. The liquid junction potential was not corrected.

Miniature IPSCs in CA2 PCs were collected using Minianalysis (version 6.0.7). All detected events were visually examined and subsequently validated by the user. Traces in the figures were digitally low pass filtered with a Gaussian filter at 1 kHz. Cumulative distributions typically contained ∼200 events. Further analysis was performed in clampFit 10.4 (Molecular devices), Stimfit 0.14 (Christoph Schmidt-Hieber, Institut Pasteur, https://github.com/neurodroid/stimfit^49^), Excel (Microsoft) and Mathematica 13 (Wolfram Research, Champaign, IL).

### Quantification and statistical analysis

All values were reported as mean ± SEM. Normality of the data was tested using a Shapiro– Wilk test and parametric or non-parametric tests were performed accordingly. Details for the statistical tests, n numbers and p-values are given in the figure legends. For post-hoc analysis of two-way ANOVA, Tukey’s multiple comparison test was performed. Differences with p < 0.05 were considered significant.

## Figure legends

***Figure S1. No change in total number of hippocampal GABAergic interneurons at three month upon loss of Ccp1***

**(A)** Representative images of hippocampi from 3 months old WT and Ccp1 cKO mice immunolabled to detect GFP (green), nuclei were counterstained with Dapi (blue). WT and Ccp1 cKO animals were crossed with ROSA-YFP to permanently stain interneurons. (**B**) Quantification of interneuron density. The total number of GFP+ interneurons was divided by the area of CA1, CA2 and CA3. WT = 3 and Ccp1 cKO = 4, a Mann-Whitney test was performed (p=0.2286).

***Figure S2. Interneuron protein glutamylation levels are consistent across hippocampal regions***

**A)** Representative close up images of hippocampi immunolabeled with RGS14 (cyan), PolyE (red), and PV (green) in WT and Ccp1 cKO three months old mice. PolyE^+^ labeling increased in PV^+^ interneurons from Ccpi CKO as compared to WT mice hippocampi (compare white arrowheads). (**B**) Histogram of the average PolyE intensity in PV^+^ interneurons across hippocampal regions from WT (green bars) and Ccp1 cKO (red bars) mice. PolyE staining intensity was normalized to the intensity of the surrounding hippocampal tissue. Each dot on the graph represents the average PolyE intensity for all PV^+^ interneurons analyzed in one animal. WT: n=5 CCP1cKO n= 5. A two-way ANOVA was conducted after confirmation of normal data distribution, * = p < 0.5, ** = p < 0.1.

***Figure S3. Characterization of DIV15 hippocampal cultures***

**(A)** Immunolabeling of DIV 15 hippocampal culture isolated from E17.5 WT (Dlx5,6Cre-GFP) embryos, showing GFP (green), PV (red) and NeuN (light grey) and merge. Grenn arrows point interneurons (GFP^+^; NeuN^+^), yellow arrows point PV^+^ interneurons (GFP^+^; PV^+^, NeuN^+^), white arrows point projection neurons (NeuN^+^). Nuclei counterstaining by Dapi staining (blue)(**B**-**C**) Proportion of WT and Ccp1 cKO interneurons assessed as the ratio between GFP^+^/NeuN^+^ and GFP^-^/NeuN^+^ neurons. Data were collected from 3 independent experiments per genotype. (**D**-**E**) Proportion of PV^+^ interneurons in WT and Ccp1 cKO assessed as the ratio between PV^+^/GFP^+^ and PV^-^/GFP^+^ neurons. Data were collected from 3 independent experiments per genotype. (**F**) Immunolabeling of a WT and Ccp1 cKO DIV15 interneurons showing GFP (green), PolyE (red) and PV (light grey) or merge stainings. The Ccp1 cKO interneuron shows accumulation of hyperglutamylation signal.

***Figure S4. Workflow for MT-dependent transport analysis in hippocampal interneurons***

**(A)** Glass-bottom dishes with a gridded coverslip were used to seed dissociated hippocampi of E17.5 WT or Ccp1 cKO embryos that were cultured for 15 days. (**B**) At DIV15 GFP^+^ interneurons were randomly selected and their position on the grid was recorded after axonal transport recording, as exemplified by red numbers. (**C**) Interneurons were labelled with LysoTracker and their MT-dependent transport was recorded along the proximal neurite for each neuron. (**D**) Neurons were fixed and stained for GFP and PV after the recording session. Using the annotated positions on the grid, each movie was retrospectively assigned to its corresponding neuron and sorted based on PV expression. (**E**-**K**) Histograms of MT-dependent transport of lysosomes in GFP^+^,PV^-^ interneurons showing the anterograde velocity in both genotypes. The fraction of movement when lysosomes travel away from the cell body is considered. The number of lysosomes with anterograde movement in this analysis is 363 for WT and 519 for Ccp1 cKO. A Mann-Whitney test was conducted (p = 0.2017) (**E**). Histogram of retrograde velocity in both genotypes. Like anterograde velocity, retrograde velocity only considers the fraction of movement when lysosomes travel towards the cell body. The number of lysosomes with retrograde movement in this analysis is 522 for WT and 604 for Ccp1 cKO. A Mann-Whitney test was conducted (p = 0.0699) (**F**). Histogram of the fraction of time spent moving in both genotypes. This analysis includes all lysosomes tracked, including motile and stationary ones and it measures the percentage of time in movement for each lysosome. A threshold of 0.1 µm/s was established below which vesicles are considered immotile. All lysosomes that were tracked were included in this analysis for a total of 1303 and 1506 in WT and Ccp1 cKO, respectively. A Mann-Whitney test was conducted (p < 0.1107) (**G**). Histogram of the fraction of time pausing for lysosmes in both genotypes. This measurement only considers those that were motile for at least a portion of the movie and represents the percentage of time spent with a speed < 0.1 µm/s, which is considered as pausing. A total of 689 and 833 motile lysosomes were included for the WT and Ccp1 cKO conditions, respectively. A Mann-Whitney test was conducted (p < 0.0001) (**H**). Histogram of the percentage of immotile vesicles. Lysosomes are considered immotile if their instantaneous speed is < 0.1µm/s for the whole duration of the movie. For this analysis, the proportion of immotile lysosomes in each neuron was calculated. Here, 39 and 47 PV^+^ interneurons were recorded in the WT and Ccp1 cKO conditions respectively, from 3 separate experiments. An unpaired t-student test was conducted (p = 0.7165) (**I**). (**J**-**K**) Kymographs represent the tracks that were manually traced where blue lines correspond to time pausing, green lines anterograde movement and red lines retrograde movement in PV^-^ interneurons from both genotypes.

***Figure S5. Hippocampal inhibitory synapses and electrophysiological properties of CA2 PCs***

(**A**-**H**) Quantification of inhibitory synapses in the CA1, CA2 and CA3 regions of the hippocampus subdivided in stratum oriens, pyramidale, and radiatum/lacunosum in six WT and six Ccp1 cKO 3 months old mice. Quantification was performed by assessing the density of colocalizing VGAT; gephyrin punctae. Unpaired t-student test was conducted, p-values are p=0.1429 (**A**), p=0.5655 (**B**), p=0.0769 (**C**), p=0.0428 (**D**), p=0.0650 (**E**), p=0.4545 (**F**), p=0.2719 (G), 0.2958 (H). (**I**, **L**) Representative traces of sIPSCs in a CA2 PC at -70 mV in WT slices (**I**, green trace) and Ccp1 cKO slices (**L**, red trace). 20 mM D-AP5, and 10 mM CNQX were added to external solution to block excitatory synaptic activity. Inset (**I**) shows average sIPSC at expanded timescale (average from 2,740 [9 WT neurons] and inset (**L**) average from 2,341 [8 cKO neurons] single events). (**J**) Cumulative histograms of sIPSC amplitude (light green: from 9 individual PCs of WT mice; thick green: average of WT data; light red: from 8 individual PCs of Ccp1 cKO mice; red: average of Ccp1 cKO data). (**K**) Boxplots of sIPSC peak amplitude. Data from 9 WT neurons and 8 cKO neurons. Averaged sIPSC amplitude was slightly but not significantly larger in neurons from Ccp1 cKO neurons in comparison to WT neurons (t test, p=0.0963). (**M**) Cumulative histograms of sIPSC inter-event interval (light green: from 9 individual PCs of WT mice; thick green: average of WT data; light red: from 8 individual PCs of Ccp1 cKO mice; red: average of Ccp1 cKO data). (**N**) Boxplots of sIPSC frequency. Data from 9 WT neurons and 8 cKO neurons, respectively. Averaged sIPSC frequency was not changed in cKO neurons when comparing to control neurons (t test, p=0.97). In box plots (**K** and **N**), horizontal lines represent median; boxes, quartiles; whiskers, most extreme data points; and single points, data from individual experiments. (**O**-**P**) Voltage traces recorded in whole-cell current-clamp in a CA2 PC in WT (**O**) and Ccp1 cKO (**P**) mouse slice in response to a 1 s long hyperpolarizing (-150 pA) and depolarizing (150 pA) current injections. (**Q**-**R**) resting membrane potential (**Q**) and input resistance (**R**) in CA2 PCs. Data are from 9 PCs in WT and 11 PCs in Ccp1 cKO. The resting membrane potential was in tendency lower in Ccp1 cKO neurons in comparison to WT neurons (Mann-Whitney test, p=0.0615) and the input resistance was slightly higher in Ccp1 cKO neurons in comparison to WT neurons (t test, p=0.0501). In box plots (**Q**-**R**), horizontal lines represent median; boxes, quartiles; whiskers, most extreme data points; and single points, data from individual experiments. (**S**) Input-output curve showing average action potential number of PCs in response to current injections of increasing amplitudes (0-400 pA) in WT versus Ccp1 cKO. Current to action potential number plots for CA2 PCs shows a slight non-significant increase of action potential number for neurons from Ccp1 cKO mice (two-way RM ANOVA, current step: F_(8,136)_ = 46.94; p < 0.0001; genotype: F_(1,17)_ = 1.961; p = 0.1794; interaction: F_(8,136)_ = 1.034; p = 0.4134; n=9 for WT and n=11 for Ccp1 cKO). Data points in (**S**) represent mean ± SEM.

***Figure S6. Analysis of inhibitory synapses in the hippocampal CA1, CA2 and CA3 regions***

Tile-scan images of synapse settings were captured in portions of the CA1, CA2 and CA3 fields (Yellow dotted lines). The CA1/CA2 boundary was determined based on DAPI staining as shown in panel. CA1, CA2 and CA3 subregions were then divided in stratum oriens (SO), stratum pyramidale (SP) and stratum radiatum/lacunosum (SRL) for synapse analysis.

***Figure S7. Single spot detection to quantify inhibitory hippocampal synapses***

**(A)** Magnification of inhibitory synapses stained with VGAT and Gephyrin in the hippocampus of WT mice at 3 months. (**B**) Spots were created based on VGAT and gephyrin signal with the Imaris software. Green and red spots are generated based on VGAT and gephyrin staining respectively. Yellow spots correspond to gephyrin spots that are within 400nm of a VGAT spot. The number of yellow spots was considered as the number of inhibitory synapses and biologically corresponds to the number of post-synaptic sites that co-distribute with pre-synaptic sites. (**C**) Merged image representing the accurate detection of synapses using the Imaris method of quantification.

## Acknowledgements

D.E., B.L., and L.N. are Research Associates and Research Director from F.R.S-F.N.R.S., respectively. We thank the members of the Nguyen lab for the discussion and feedback on the present work. We thank the GIGA imaging platform and particularly Alexandre Hego for helping with the acquisition of confocal images. We would further like to thank Laura Lebrun and Veronique Henriot (Insitut Curie) for technical assistance.

The work performed in the Nguyen laboratory is supported by ULiège (Crédit Classique), the F.R.S.-F.N.R.S. (PDR T.0185.20; EOS 0019118F-RG36), the WEL Research Institute (CR-2022A-12), the Fonds Leon Fredericq, the Fondation Simone et Pierre Clerdent, the Fondation Médicale Reine Elisabeth, the ERANET Neuron (STEM-MCD and NeuroTalk), the Win2Wal (ChipOmics; #2010126), and the ERC-Synergy (UNFOLD). The Janke lab is supported by the French National Research Agency (ANR) awards ANR-20-CE13-0011 and the Fondation pour la Recherche Medicale (FRM) grant MND202003011485.

## Bibliography

1. Dent, E.W. (2020). Dynamic microtubules at the synapse. Curr Opin Neurobiol 63, 9–14. 10.1016/j.conb.2020.01.003.

2. Guedes-Dias, P., and Holzbaur, E.L.F. (2019). Axonal transport: Driving synaptic function. Science 366. 10.1126/science.aaw9997.

3. Roll-Mecak, A. (2019). How cells exploit tubulin diversity to build functional cellular microtubule mosaics. Curr Opin Cell Biol 56, 102–108. 10.1016/j.ceb.2018.10.009.

4. Janke, C., and Magiera, M.M. (2020). The tubulin code and its role in controlling microtubule properties and functions. Nat Rev Mol Cell Biol 21, 307–326. 10.1038/s41580-020-0214-3.

5. van Dijk, J., Miro, J., Strub, J.M., Lacroix, B., van Dorsselaer, A., Edde, B., and Janke, C. (2008). Polyglutamylation is a post-translational modification with a broad range of substrates. J Biol Chem 283, 3915–3922. 10.1074/jbc.M705813200.

6. Chen, J., Zehr, E.A., Gruschus, J.M., Szyk, A., Liu, Y., Tanner, M.E., Tjandra, N., and Roll-Mecak, A. (2024). Tubulin code eraser CCP5 binds branch glutamates by substrate deformation. Nature 631, 905–912. 10.1038/s41586-024-07699-0.

7. Rogowski, K., van Dijk, J., Magiera, M.M., Bosc, C., Deloulme, J.C., Bosson, A., Peris, L., Gold, N.D., Lacroix, B., Bosch Grau, M., et al. (2010). A family of protein-deglutamylating enzymes associated with neurodegeneration. Cell 143, 564–578. 10.1016/j.cell.2010.10.014.

8. Magiera, M.M., Bodakuntla, S., Ziak, J., Lacomme, S., Marques Sousa, P., Leboucher, S., Hausrat, T.J., Bosc, C., Andrieux, A., Kneussel, M., et al. (2018). Excessive tubulin polyglutamylation causes neurodegeneration and perturbs neuronal transport. EMBO J 37. 10.15252/embj.2018100440.

9. Rogowski, K., van Dijk, J., Magiera, M.M., Bosc, C., Deloulme, J.C., Bosson, A., Peris, L., Gold, N.D., Lacroix, B., Grau, M.B., et al. (2010). A Family of Protein-Deglutamylating Enzymes Associated with Neurodegeneration. Cell 143, 564–578. 10.1016/j.cell.2010.10.014.

10. Bodakuntla, S., Schnitzler, A., Villablanca, C., Gonzalez-Billault, C., Bieche, I., Janke, C., and Magiera, M.M. (2020). Tubulin polyglutamylation is a general traffic-control mechanism in hippocampal neurons. J Cell Sci 133. 10.1242/jcs.241802.

11. Gilmore-Hall, S., Kuo, J., Ward, J.M., Zahra, R., Morrison, R.S., Perkins, G., and La Spada, A.R. (2019). CCP1 promotes mitochondrial fusion and motility to prevent Purkinje cell neuron loss in pcd mice. J Cell Biol 218, 206–219. 10.1083/jcb.201709028.

12. Bodakuntla, S., Yuan, X., Genova, M., Gadadhar, S., Leboucher, S., Birling, M.C., Klein, D., Martini, R., Janke, C., and Magiera, M.M. (2021). Distinct roles of alpha- and beta-tubulin polyglutamylation in controlling axonal transport and in neurodegeneration. EMBO J 40, e108498. 10.15252/embj.2021108498.

13. Silva, C.G., Peyre, E., Adhikari, M.H., Tielens, S., Tanco, S., Van Damme, P., Magno, L., Krusy, N., Agirman, G., Magiera, M.M., et al. (2018). Cell-Intrinsic Control of Interneuron Migration Drives Cortical Morphogenesis. Cell 172, 1063–1078 e1019. 10.1016/j.cell.2018.01.031.

14. Butt, S.J., Fuccillo, M., Nery, S., Noctor, S., Kriegstein, A., Corbin, J.G., and Fishell, G. (2005). The temporal and spatial origins of cortical interneurons predict their physiological subtype. Neuron 48, 591–604. 10.1016/j.neuron.2005.09.034.

15. Tricoire, L., Pelkey, K.A., Erkkila, B.E., Jeffries, B.W., Yuan, X., and McBain, C.J. (2011). A blueprint for the spatiotemporal origins of mouse hippocampal interneuron diversity. J Neurosci 31, 10948–10970. 10.1523/JNEUROSCI.0323-11.2011.

16. Booker, S.A., and Vida, I. (2018). Morphological diversity and connectivity of hippocampal interneurons. Cell Tissue Res 373, 619–641. 10.1007/s00441-018-2882-2.

17. Stenman, J., Toresson, H., and Campbell, K. (2003). Identification of two distinct progenitor populations in the lateral ganglionic eminence: implications for striatal and olfactory bulb neurogenesis. J Neurosci 23, 167–174.

18. Berezniuk, I., Sironi, J., Callaway, M.B., Castro, L.M., Hirata, I.Y., Ferro, E.S., and Fricker, L.D. (2010). CCP1/Nna1 functions in protein turnover in mouse brain: Implications for cell death in Purkinje cell degeneration mice. FASEB J 24, 1813–1823. 10.1096/fj.09-147942.

19. Shang, Y., Li, B., and Gorovsky, M.A. (2002). Tetrahymena thermophila contains a conventional gamma-tubulin that is differentially required for the maintenance of different microtubule-organizing centers. J Cell Biol 158, 1195–1206. 10.1083/jcb.200205101.

20. Evans, P.R., Parra-Bueno, P., Smirnov, M.S., Lustberg, D.J., Dudek, S.M., Hepler, J.R., and Yasuda, R. (2018). RGS14 Restricts Plasticity in Hippocampal CA2 by Limiting Postsynaptic Calcium Signaling. eNeuro 5. 10.1523/ENEURO.0353-17.2018.

21. Chaudhry, F.A., Reimer, R.J., Bellocchio, E.E., Danbolt, N.C., Osen, K.K., Edwards, R.H., and Storm-Mathisen, J. (1998). The vesicular GABA transporter, VGAT, localizes to synaptic vesicles in sets of glycinergic as well as GABAergic neurons. J Neurosci 18, 9733–9750. 10.1523/JNEUROSCI.18-23-09733.1998.

22. Kneussel, M., Brandstatter, J.H., Gasnier, B., Feng, G., Sanes, J.R., and Betz, H. (2001). Gephyrin-independent clustering of postsynaptic GABA(A) receptor subtypes. Mol Cell Neurosci 17, 973–982. 10.1006/mcne.2001.0983.

23. Dobie, F.A., and Craig, A.M. (2011). Inhibitory synapse dynamics: coordinated presynaptic and postsynaptic mobility and the major contribution of recycled vesicles to new synapse formation. J Neurosci 31, 10481–10493. 10.1523/JNEUROSCI.6023-10.2011.

24. Pawelzik, H., Hughes, D.I., and Thomson, A.M. (2002). Physiological and morphological diversity of immunocytochemically defined parvalbumin- and cholecystokinin-positive interneurones in CA1 of the adult rat hippocampus. J Comp Neurol 443, 346–367. 10.1002/cne.10118.

25. Frerking, M., Borges, S., and Wilson, M. (1995). Variation in GABA mini amplitude is the consequence of variation in transmitter concentration. Neuron 15, 885–895. 10.1016/0896-6273(95)90179-5.

26. Freund, T.F., and Katona, I. (2007). Perisomatic inhibition. Neuron 56, 33–42. 10.1016/j.neuron.2007.09.012.

27. Mercer, A., Trigg, H.L., and Thomson, A.M. (2007). Characterization of neurons in the CA2 subfield of the adult rat hippocampus. J Neurosci 27, 7329–7338. 10.1523/JNEUROSCI.1829-07.2007.

28. Engel, D., Pahner, I., Schulze, K., Frahm, C., Jarry, H., Ahnert-Hilger, G., and Draguhn, A. (2001). Plasticity of rat central inhibitory synapses through GABA metabolism. The Journal of physiology 535, 473–482. 10.1111/j.1469-7793.2001.00473.x.

29. Soltesz, I., Smetters, D.K., and Mody, I. (1995). Tonic inhibition originates from synapses close to the soma. Neuron 14, 1273–1283. 10.1016/0896-6273(95)90274-0.

30. Nevian, T., Larkum, M.E., Polsky, A., and Schiller, J. (2007). Properties of basal dendrites of layer 5 pyramidal neurons: a direct patch-clamp recording study. Nat Neurosci 10, 206–214. 10.1038/nn1826.

31. Shao, L.R., and Dudek, F.E. (2005). Changes in mIPSCs and sIPSCs after kainate treatment: evidence for loss of inhibitory input to dentate granule cells and possible compensatory responses. Journal of neurophysiology 94, 952–960. 10.1152/jn.01342.2004.

32. Ikegami, K., Heier, R.L., Taruishi, M., Takagi, H., Mukai, M., Shimma, S., Taira, S., Hatanaka, K., Morone, N., Yao, I., et al. (2007). Loss of alpha-tubulin polyglutamylation in ROSA22 mice is associated with abnormal targeting of KIF1A and modulated synaptic function. Proc Natl Acad Sci U S A 104, 3213–3218. 10.1073/pnas.0611547104.

33. Lopes, A.T., Hausrat, T.J., Heisler, F.F., Gromova, K.V., Lombino, F.L., Fischer, T., Ruschkies, L., Breiden, P., Thies, E., Hermans-Borgmeyer, I., et al. (2020). Spastin depletion increases tubulin polyglutamylation and impairs kinesin-mediated neuronal transport, leading to working and associative memory deficits. PLoS Biol 18, e3000820. 10.1371/journal.pbio.3000820.

34. Millecamps, S., and Julien, J.P. (2013). Axonal transport deficits and neurodegenerative diseases. Nat Rev Neurosci 14, 161–176. 10.1038/nrn3380.

35. Chevaleyre, V., and Siegelbaum, S.A. (2010). Strong CA2 pyramidal neuron synapses define a powerful disynaptic cortico-hippocampal loop. Neuron 66, 560–572. 10.1016/j.neuron.2010.04.013.

36. Piskorowski, R.A., and Chevaleyre, V. (2023). Hippocampal area CA2: interneuron disfunction during pathological states. Front Neural Circuits 17, 1181032. 10.3389/fncir.2023.1181032.

37. Bateup, H.S., Johnson, C.A., Denefrio, C.L., Saulnier, J.L., Kornacker, K., and Sabatini, B.L. (2013). Excitatory/inhibitory synaptic imbalance leads to hippocampal hyperexcitability in mouse models of tuberous sclerosis. Neuron 78, 510–522. 10.1016/j.neuron.2013.03.017.

38. Gibson, J.R., Bartley, A.F., Hays, S.A., and Huber, K.M. (2008). Imbalance of neocortical excitation and inhibition and altered UP states reflect network hyperexcitability in the mouse model of fragile X syndrome. Journal of neurophysiology 100, 2615–2626. 10.1152/jn.90752.2008.

39. Banerjee, A., Rikhye, R.V., Breton-Provencher, V., Tang, X., Li, C., Li, K., Runyan, C.A., Fu, Z., Jaenisch, R., and Sur, M. (2016). Jointly reduced inhibition and excitation underlies circuit-wide changes in cortical processing in Rett syndrome. Proc Natl Acad Sci U S A 113, E7287–E7296. 10.1073/pnas.1615330113.

40. Hitti, F.L., and Siegelbaum, S.A. (2014). The hippocampal CA2 region is essential for social memory. Nature 508, 88–92. 10.1038/nature13028.

41. Bischofberger, J., Engel, D., Frotscher, M., and Jonas, P. (2006). Timing and efficacy of transmitter release at mossy fiber synapses in the hippocampal network. Pflugers Archiv: European journal of physiology 453, 361–372. 10.1007/s00424-006-0093-2.

42. Piskorowski, R.A., and Chevaleyre, V. (2013). Delta-opioid receptors mediate unique plasticity onto parvalbumin-expressing interneurons in area CA2 of the hippocampus. J Neurosci 33, 14567–14578. 10.1523/JNEUROSCI.0649-13.2013.

43. Schneider Gasser, E.M., Straub, C.J., Panzanelli, P., Weinmann, O., Sassoe-Pognetto, M., and Fritschy, J.M. (2006). Immunofluorescence in brain sections: simultaneous detection of presynaptic and postsynaptic proteins in identified neurons. Nat Protoc 1, 1887–1897. 10.1038/nprot.2006.265.

44. Falzone, T.L., and Stokin, G.B. (2012). Imaging amyloid precursor protein in vivo: an axonal transport assay. Methods Mol Biol 846, 295–303. 10.1007/978-1-61779-536-7_25.

45. Zala, D., Hinckelmann, M.V., Yu, H., Lyra da Cunha, M.M., Liot, G., Cordelieres, F.P., Marco, S., and Saudou, F. (2013). Vesicular glycolysis provides on-board energy for fast axonal transport. Cell 152, 479–491. 10.1016/j.cell.2012.12.029.

46. Bischofberger, J., Engel, D., Li, L., Geiger, J.R., and Jonas, P. (2006). Patch-clamp recording from mossy fiber terminals in hippocampal slices. Nat Protoc 1, 2075–2081. 10.1038/nprot.2006.312.

47. Engel, D., and Jonas, P. (2005). Presynaptic action potential amplification by voltage-gated Na+ channels in hippocampal mossy fiber boutons. Neuron 45, 405–417. 10.1016/j.neuron.2004.12.048.

48. Fernandez-Lamo, I., Gomez-Dominguez, D., Sanchez-Aguilera, A., Oliva, A., Morales, A.V., Valero, M., Cid, E., Berenyi, A., and Menendez de la Prida, L. (2019). Proximodistal Organization of the CA2 Hippocampal Area. Cell Rep 26, 1734–1746 e1736. 10.1016/j.celrep.2019.01.060.

49. Guzman, S.J., Schlogl, A., and Schmidt-Hieber, C. (2014). Stimfit: quantifying electrophysiological data with Python. Front Neuroinform 8, 16. 10.3389/fninf.2014.00016.

