## supplemental figures for "Ccp1 depletion disrupts network integration of hippocampal parvalbumin interneurons"

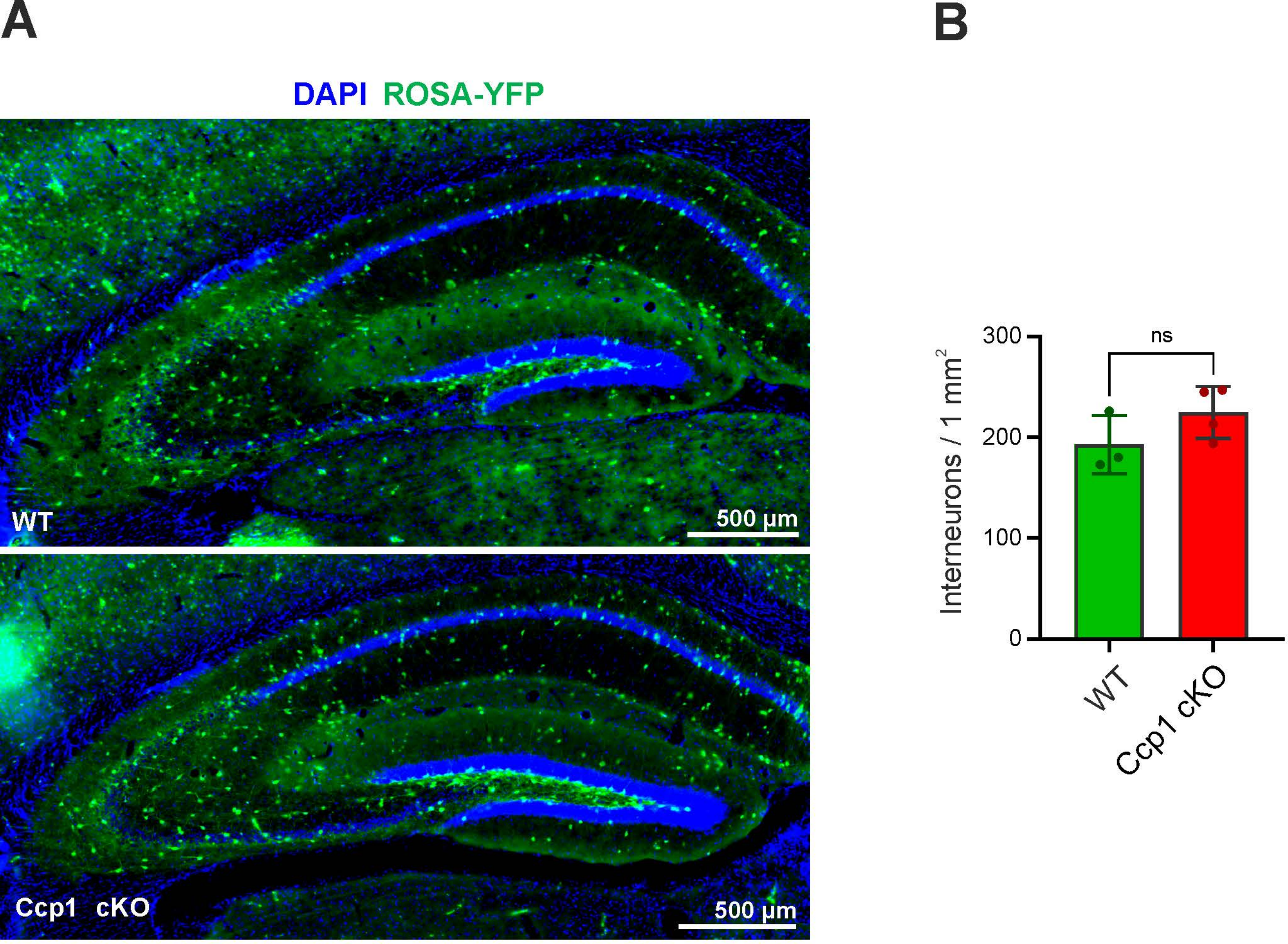
A
DAPI ROSA-YFP
WT
500 µm
Ccp1 cKO
500 µm
B
ns
Interneurons / 1 mm2
300
200
100
0
WT
Ccp1 cKO

Figure S1

A

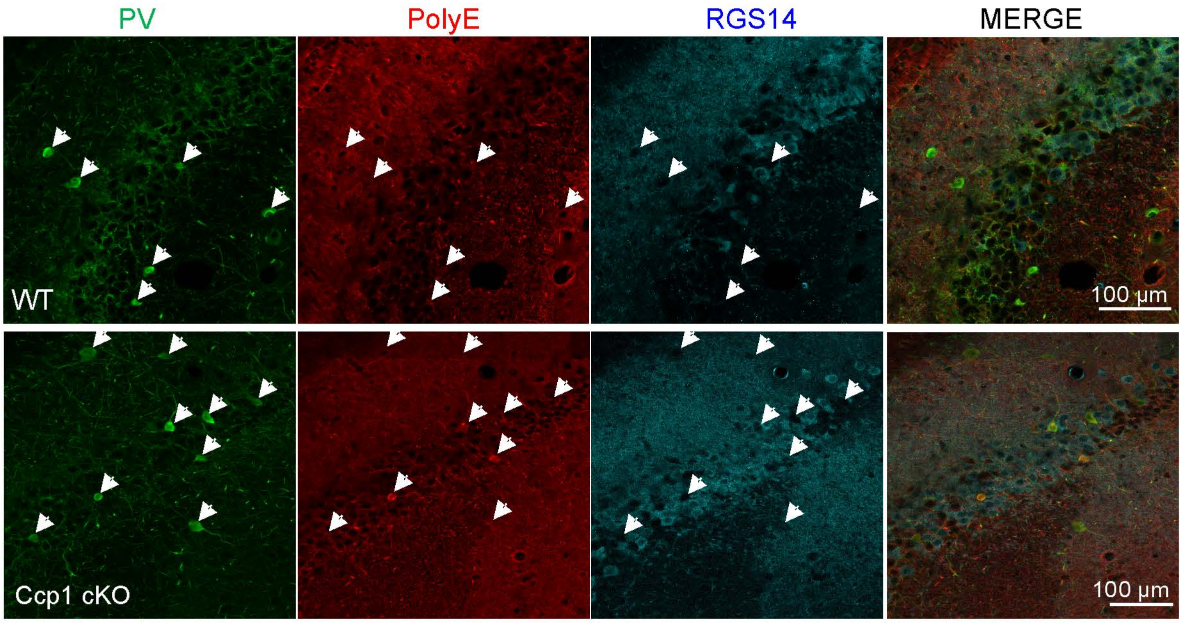
PV
PolyE
RGS14
MERGE
WT
Ccp1 cKO
100 µm
100 µm

B

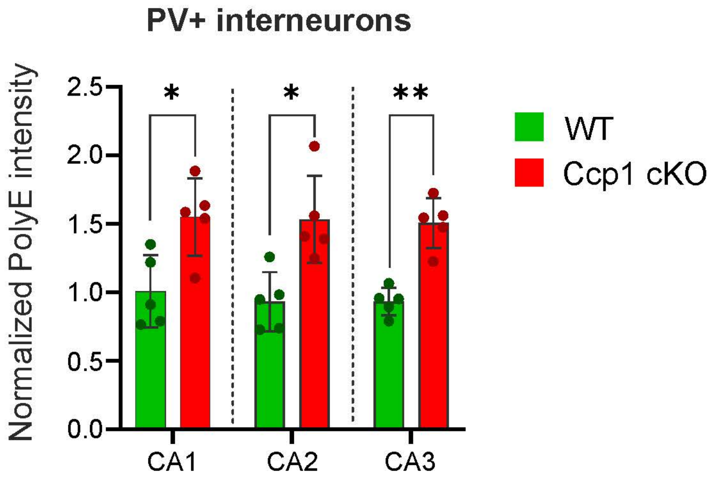
PV+ interneurons
Normalized PolyE intensity
2.5
2.0
1.5
1.0
0.5
0.0
*
*
**
CA1
CA2
CA3
WT
Ccp1 cKO

Figure S2

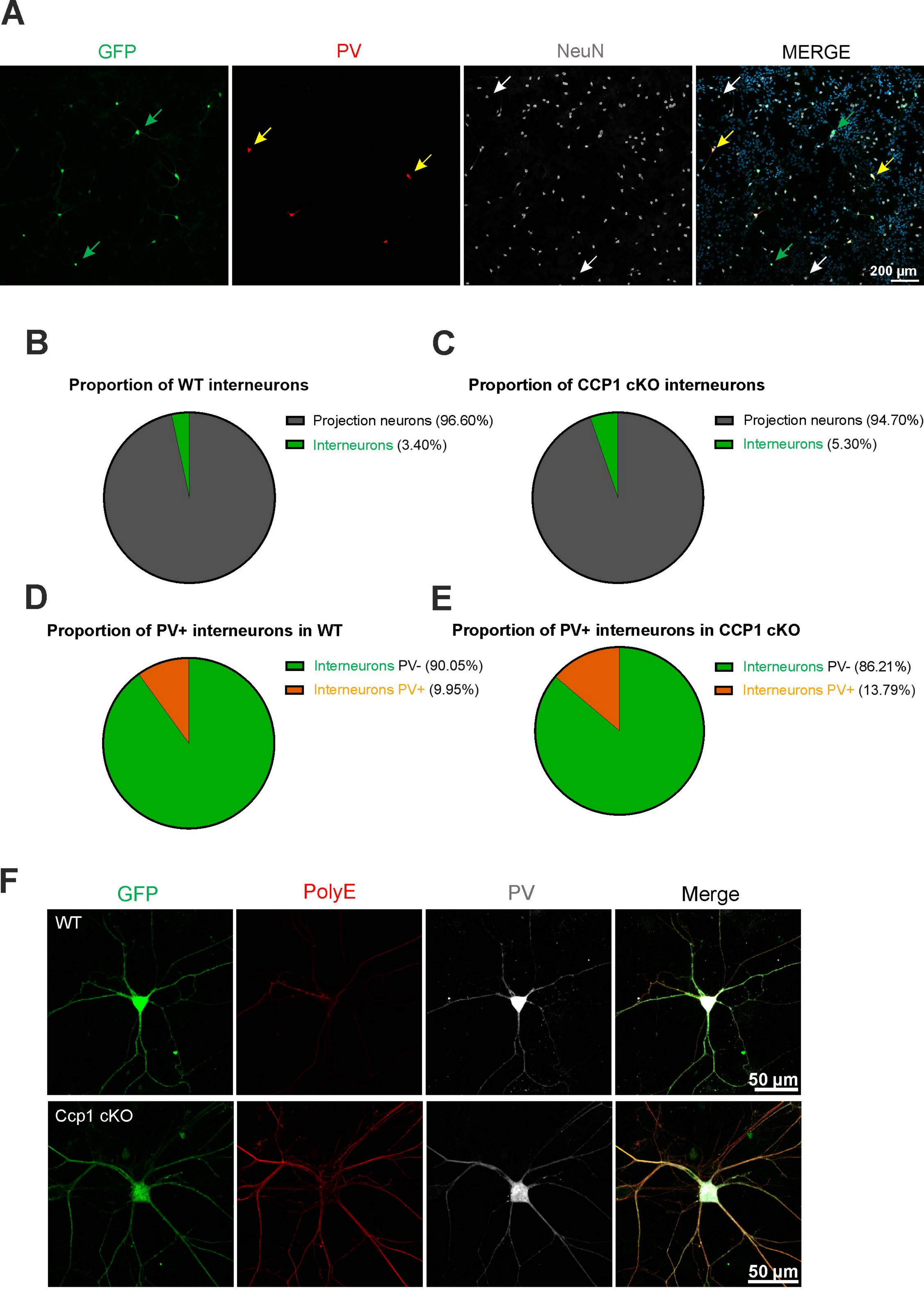
A
GFP
PV
NeuN
MERGE
200 µm
B
Proportion of WT interneurons
Projection neurons (96.60%)
Interneurons (3.40%)
C
Proportion of CCP1 cKO interneurons
Projection neurons (94.70%)
Interneurons (5.30%)
D
Proportion of PV+ interneurons in WT
Interneurons PV- (90.05%)
Interneurons PV+ (9.95%)
E
Proportion of PV+ interneurons in CCP1 cKO
Interneurons PV- (86.21%)
Interneurons PV+ (13.79%)
F
GFP
PolyE
PV
Merge
WT
50 µm
Ccp1 cKO
50 µm

Figure S3

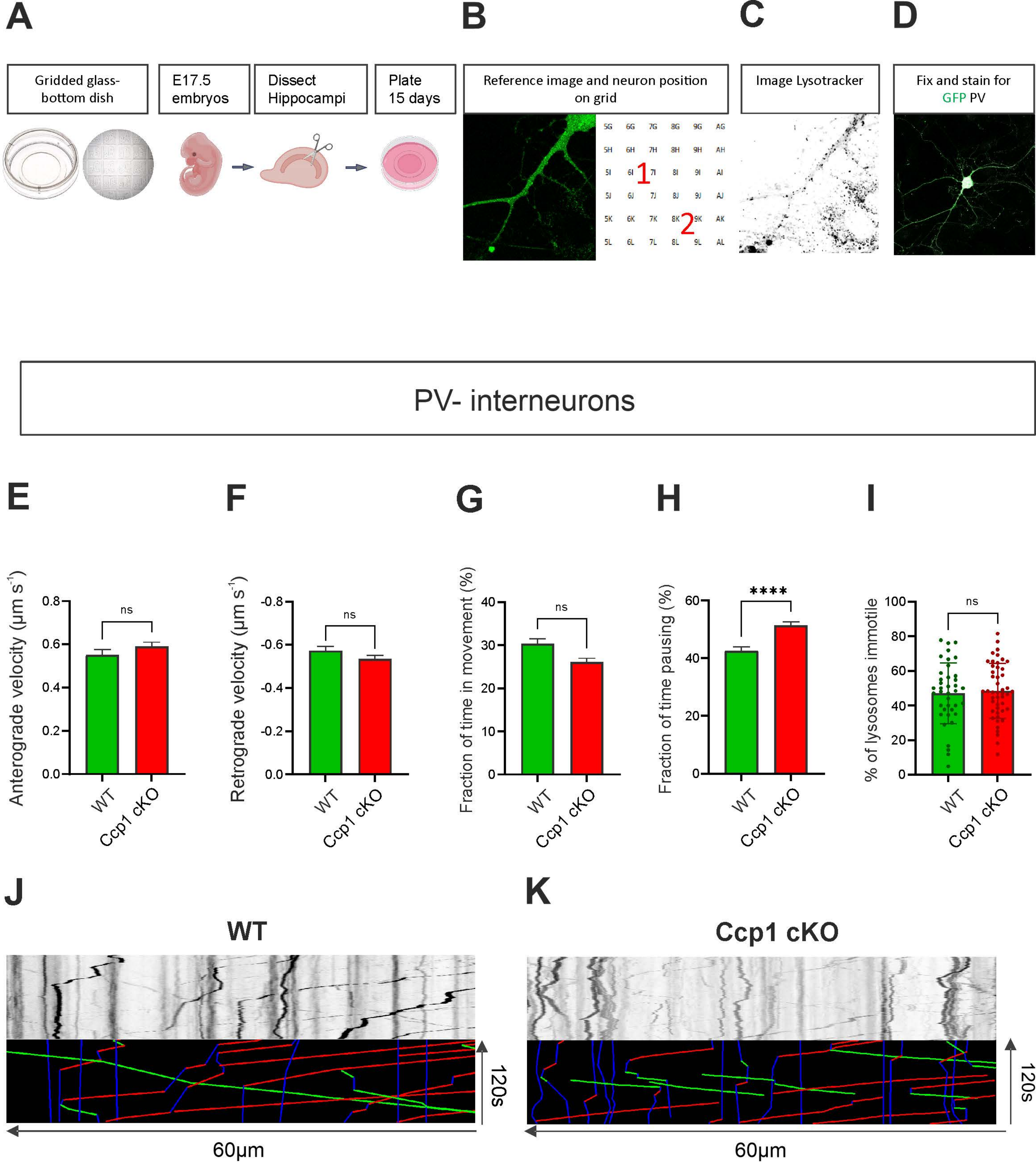

A
B
C
D
Gridded glass-bottom dish
E17.5 embryos
Dissect Hippocampi
Plate 15 days
Reference image and neuron position on grid
Image Lysotracker
Fix and stain for GFP PV
PV- interneurons
E
F
G
H
I
Anterograde velocity (μm s⁻¹)
Retrograde velocity (μm s⁻¹)
Fraction of time in movement (%)
Fraction of time pausing (%)
% of lysosomes immotile
ns
****
WT
Ccp1 cKO
J
K
WT
Ccp1 cKO
120s
60μm


Figure S4

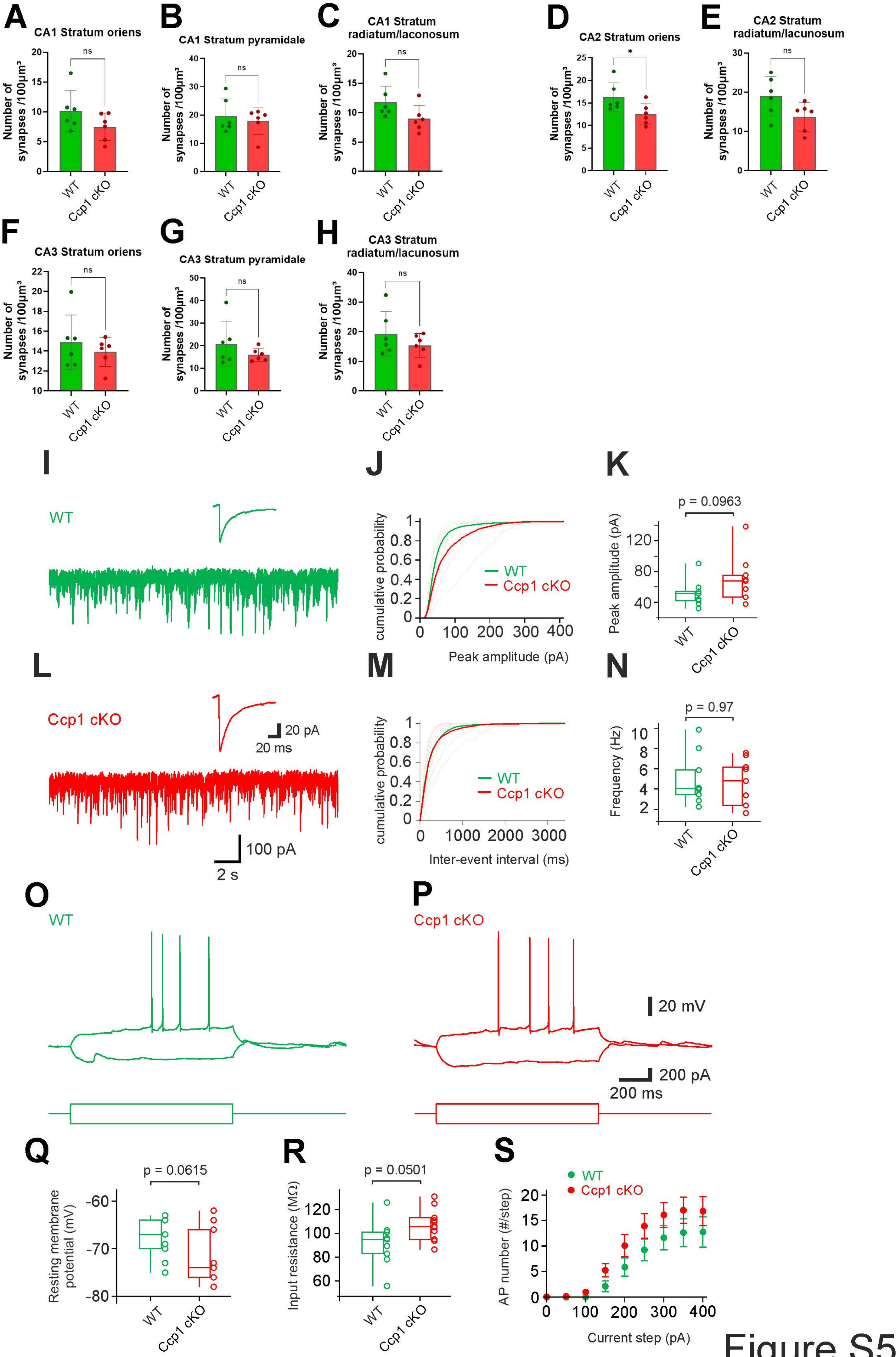

A
CA1 Stratum oriens
B
CA1 Stratum pyramidale
C
CA1 Stratum radiatum/laconosum
D
CA2 Stratum oriens
E
CA2 Stratum radiatum/lacunosum
F
CA3 Stratum oriens
G
CA3 Stratum pyramidale
H
CA3 Stratum radiatum/lacunosum
Number of synapses /100µm³
ns
*
WT
Ccp1 cKO
I
J
K
L
M
N
cumulative probability
Peak amplitude (pA)
Inter-event interval (ms)
p = 0.0963
p = 0.97
Frequency (Hz)
20 pA
20 ms
100 pA
2 s
O
P
20 mV
200 pA
200 ms
Q
R
S
p = 0.0615
p = 0.0501
Resting membrane potential (mV)
Input resistance (MΩ)
AP number (#/step)
Current step (pA)


Figure S5

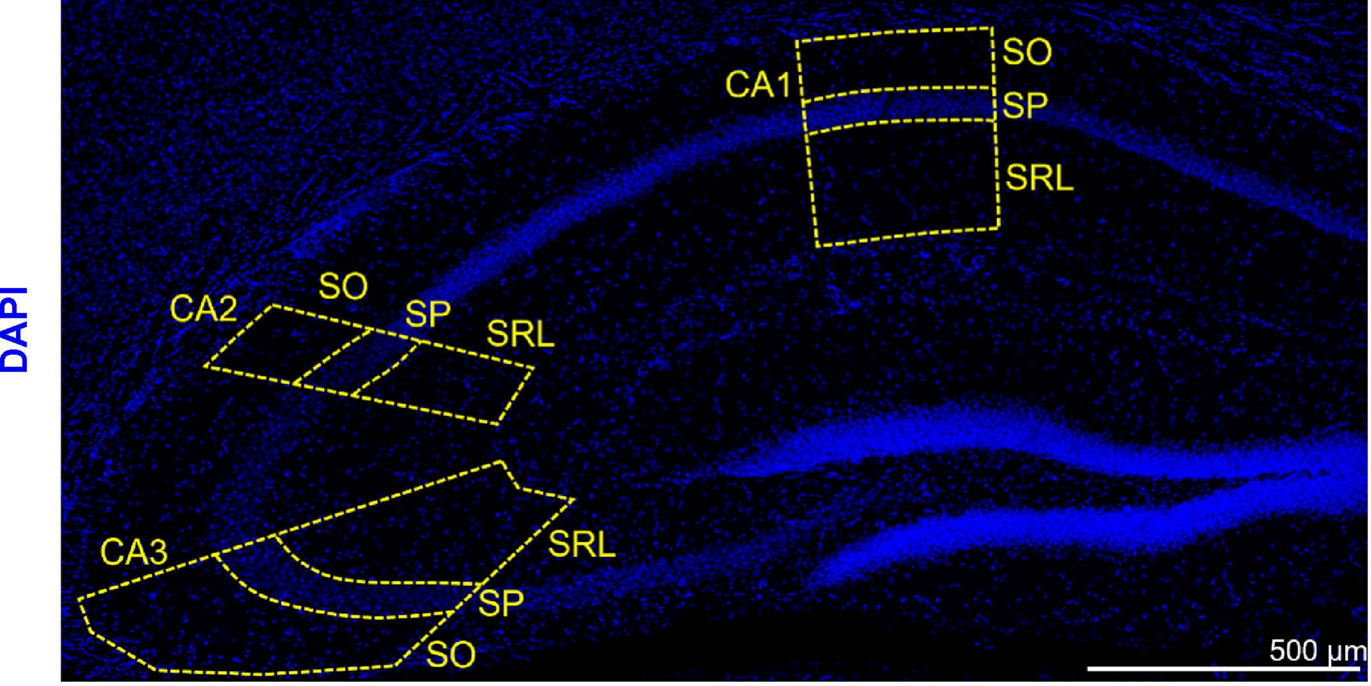


Figure S6

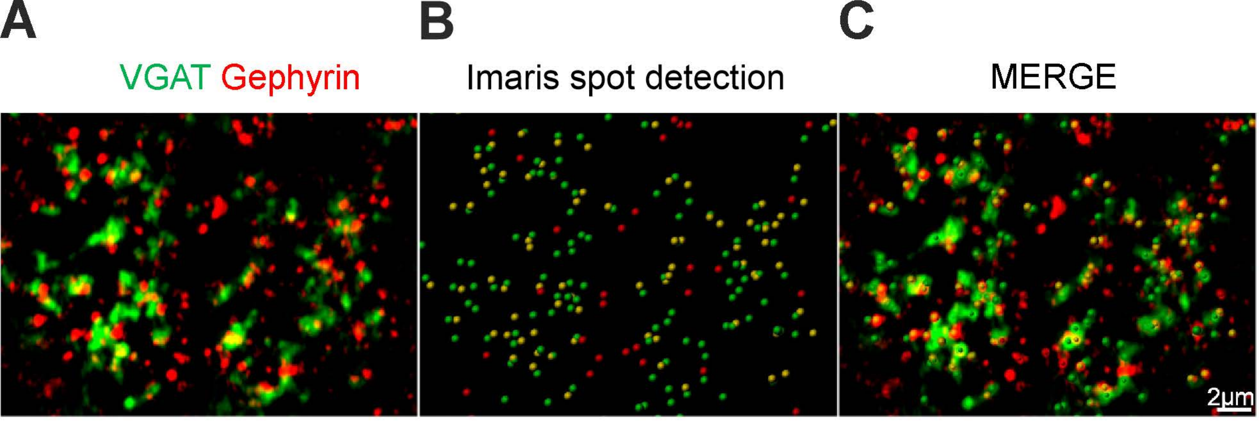


Figure S7

| Experiment | Antibodies | Host | Antigen retrieval | Dilution/ Incubation | Reference |
|---|---|---|---|---|---|
| **Figure 1** | Parvalbumin | Guinea pig | From Basescope protocol | 1:300, O/N 4°C | Synaptic systems, 106 104 |
| | Somatostatin | Rabbit | | 1:300, O/N 4°C | Peninsula laboratories, T-4102 |
| **Figure 2** | Parvalbumin | Guinea pig | DAKO, 80°C, 25min | 1:500, O/N RT | Synaptic systems, 106 104 |
| | PolyE | Rabbit | | 1:1000, O/N RT | Adipogen, AG-25B-0030 |
| | Somatostatin | Mouse | | 1:300, O/N RT | Santa Cruz, sc-55565 |
| **Figure 4A** | Parvalbumin | Guinea pig | No | 1:500, O/N 4°C | Synaptic systems, 106 104 |
| **Figure 4C** | Somatostatin | Rabbit | No | 1:500, O/N 4°C | Peninsula laboratories, T-4102 |
| **Figure 5A** | Gephyrin | Guinea Pig | See methods | 1:200, O/N 4°C | Synaptic systems, 147 318 |
| | VGAT | Chicken | | 1:500, O/N 4°C | Synaptic systems, 131 006 |
| **Figure 5C** | GFP | Rat | No | 1:1000, O/N 4°C | Nacalai tesque, 04404-26 |
| | Gephyrin | Guinea Pig | | 1:200, O/N 4°C | Synaptic systems, 147 318 |
| | VGAT | Chicken | | 1:500, O/N 4°C | Synaptic systems, 131 006 |
| **Figure 5F** | RGS14 | Mouse | No | 1:500, O/N RT | Neuromab, 75-170 |
| **Figure S1** | GFP | Goat | No | 1:500, O/N 4°C | Abcam, ab6673 |
| **Figure S2** | Parvalbumin | Guinea pig | No | 1:500, O/N RT | Synaptic systems, 106 104 |
| | PolyE | Rabbit | | 1:1000, O/N RT | Adipogen, AG-25B-0030 |
| | RGS14 | Mouse | | 1:500, O/N RT | Neuromab, 75-170 |
| **Figure S3A** | GFP | Rat | No | 1:1000, O/N 4°C | Nacalai tesque, 04404-26 |
| | Parvalbumin | Guinea pig | | 1:500, O/N 4°C | Synaptic systems, 106 104 |
| | NeuN | Rabbit | | 1:1500, O/N 4°C | Cell signaling, 12943 |
| **Figure S3F** | GFP | Rat | No | 1:1000, O/N 4°C | Nacalai tesque, 04404-26 |
| | PolyE | Rabbit | | 1:5000, O/N 4°C | Adipogen, AG-25B-0030 |
| | Parvalbumin | Guinea pig | | 1:500, O/N 4°C | Synaptic systems, 106 104 |

Table 1. List of primary antibodies used to perform the immunolabelings
